# Lysosomal Dysfunction–Mediated IgG Accumulation Promotes Endothelial Senescence and Lesion Progression in Cerebral Cavernous Malformations

**DOI:** 10.64898/2026.08.29.747964

**Authors:** Yingxi Yang, Yingfan Sun, Shaozhi Zhao, Qiuxia Zhou, Hui Wang, Raymond Sun, Ran Huo, Lan Dao, Zhenhuan Xu, Jiaqi Liu, Rong Grace Zhai, Yiyun Chen, Qi Zhang, Ziyuan Guo, Winson S. Ho, Jiguang Wang, Rongze O. Lu, Yong Cao

**Affiliations:** Department of Neurosurgery, Beijing Tiantan Hospital, Capital Medical University, Beijing, China; Division of Life Science, Department of Chemical and Biological Engineering, State Key Laboratory of Nervous System Disorders, The Hong Kong University of Science and Technology, Hong Kong SAR, China; Department of Neurological Surgery, University of California, San Francisco, CA, USA; Helen Diller Comprehensive Cancer Center, University of California, San Francisco, CA, USA; Department of Pediatrics, Cincinnati Children’s Hospital Medical Center, Cincinnati, OH, USA; Department of Neurology, Neuroscience Institute, University of Chicago, Chicago, IL, USA; Ultrastructural pathology department, Neuropathology Center, Beijing Neurosurgical Institute/ Beijing Tiantan Hospital, Capital Medical University, Beijing, China; SIAT-HKUST Joint Laboratory of Cell Evolution and Digital Health, HKUST Shenzhen-Hong Kong Collaborative Innovation Research Institute, Futian, Shenzhen, China; Beijing Neurosurgical Institute, Capital Medical University, Beijing, China

**Author notes:** These authors contributed equally to this work (Yingxi Yang, Yingfan Sun). Corresponding authors: Winson S. Ho MD, Department of Neurological Surgery, Helen Diller Comprehensive Cancer Center, University of California, San Francisco; San Francisco, California, USA., Jiguang Wang, PhD, Division of Life Science, Department of Chemical and Biological Engineering, State Key Laboratory of Nervous System Disorders, The Hong Kong University of Science and Technology, Hong Kong SAR, China., Rongze Olivia Lu, PhD, Department of Neurological Surgery, Helen Diller Comprehensive Cancer Center, Parker Institute for Cancer Immunotherapy, University of California, San Francisco; San Francisco, California, USA., Yong Cao, MD, Department of Neurosurgery, Beijing Tiantan Hospital, Capital Medical University; Beijing, China.

**Keywords:** Cerebral cavernous malformations, IgG, endothelial senescence, lysosomal mTORC1 signaling, anti-CD38 antibody therapy

## Abstract

Endothelial senescence is increasingly recognized as a driver of vascular pathology, while immunoglobulin G (IgG) has recently been reported to accumulate in aging tissues and induce senescence in macrophages and microglia. In cerebral cavernous malformations (CCMs), IgG accumulation has been obviously observed in CCM lesions, but the contribution of IgG to endothelial injury remains unclear. Using multi-omic profiling, endothelial models, and CCM mice, we identified IgG-secreting plasma cells enriched in lesions associated with endothelial senescence, hemorrhage, and disease severity. CCM loss-associated mTOR activation impaired lysosomal acidification and IgG processing, promoting intracellular IgG accumulation. IgG, in turn, induced NF-κB-dependent endothelial senescence. In vivo, BCMA-mediated plasma cell depletion attenuated lesion progression, whereas IgG supplementation partially restored disease severity. Anti-CD38 treatment likewise reduced IgG accumulation, endothelial senescence, hemorrhage, and lesion progression. These findings identify lysosomal dysfunction–mediated IgG as a pathogenic trigger of endothelial senescence and support targeting the plasma cell–IgG axis in CCM.

## INTRODUCTION

Endothelial senescence has emerged as an important contributor to vascular pathology across a broad range of diseases ^1–8^. Senescent endothelial cells exhibit impaired barrier function and secrete inflammatory mediators that remodel the surrounding tissue microenvironment ^9,10^. Recent studies have shown that immunoglobulin G (IgG) accumulates in aging tissues and can induce senescence-associated phenotypes in macrophages and microglia, potentially through activation of nuclear factor κB (NF-κB) signaling ^6,11,12^. However, whether IgG can directly induce endothelial senescence and the molecular mechanisms underlying this process remain unclear.

IgG is the most abundant immunoglobulin isotype in the circulation and has a prolonged serum half-life of approximately 7–23 days ^13^. Despite continuous exposure of vascular endothelial cells to high concentrations of circulating IgG, excessive intracellular IgG accumulation does not normally occur, indicating the presence of tightly regulated intracellular trafficking and recycling mechanisms. A key mediator of IgG homeostasis is the neonatal Fc receptor (FcRn), which binds IgG in a pH-dependent manner ^14,15^. FcRn binds IgG with high affinity in acidic intracellular compartments but with substantially lower affinity at less acidic or neutral extracellular pH ^14^. This pH dependence allows internalized IgG to be rescued from lysosomal degradation and recycled to the cell surface, where it dissociates from FcRn ^16^. Lysosomes are the principal degradative organelles for intracellular macromolecules, and maintenance of an acidic lysosomal lumen is essential for efficient cargo processing ^17^. Lysosomal acidification is regulated by pathways includingmechanistic target of rapamycin complex 1 (mTORC1) signaling and vacuolar H+-ATPase (V-ATPase), whereas bafilomycin A1, a V-ATPase inhibitor, suppresses lysosomal acidification ^18^. Lysosomal dysfunction is also increasingly recognized as a feature of cellular aging, and recent work has linked impaired lysosomal homeostasis and lysosomal storage disorders to age-associated disease ^19^.

Cerebral cavernous malformations (CCMs) are neurovascular lesions composed of abnormally dilated, thin-walled vascular channels ^20^. They affect up to 0.9% of the population ^21–23^ and can cause seizures, focal neurological deficits, and intracerebral hemorrhage ^24^. Surgical resection remains the definitive treatment for accessible symptomatic lesions but carries substantial risk for lesions located in deep or eloquent brain regions, and no pharmacological therapy has been established to prevent lesion progression or hemorrhage ^24,25^. Genetic studies have established that biallelic loss-of-function mutations in *CCM1*/*2*/*3* in endothelial cells, together with somatic activating mutations such as those affecting *MAP3K3*, converge on elevated MEKK3–KLF2/4 pathway and activated mTORC1 signaling, resulting in endothelial dysfunction and blood–brain barrier disruption ^26–42^.

Although endothelial dysfunction is central to CCM lesion initiation ^43–46^, accumulating evidence suggests that infiltrating immune cells also contribute to CCM pathogenesis ^47–52^. Barrier disruption, hemorrhage, and immune infiltration may expose CCM lesions to both circulating and locally produced immune cells and antibodies. Diverse immune-cell populations, including T cells, B cells, and macrophages, have been identified within CCM lesions ^47–50^. Previous studies have demonstrated IgG deposition, in situ B-cell clonal expansion and oligoclonality, and reduced lesion hemorrhage following B-cell depletion in CCM mouse models, supporting a role for humoral immunity in disease progression ^50,53–55^. However, how IgG accumulates within CCM lesions and affects endothelial function remains poorly understood.

Using multi-omic profiling, endothelial models, and CCM mouse models, we identified IgG-secreting plasma cells enriched in lesions characterized by endothelial senescence, junctional disruption, hemorrhage, and greater disease severity. We found that CCM loss-associated mTOR activation impaired lysosomal acidification and IgG processing, promoting intracellular IgG accumulation and persistence, whereas IgG induced endothelial senescence and barrier dysfunction through NF-κB signaling. mTOR inhibition, B-cell maturation antigen (BCMA)-mediated plasma cell depletion, and CD38-targeted treatment attenuated endothelial senescence and lesion progression. Using CCM endothelial and mice models, these findings identify lysosomal dysfunction–mediated IgG as a trigger of endothelial senescence and provide a mechanistic rationale for targeting the plasma cell–IgG axis in CCM.

## METHODS

Detailed methods are provided in the supplemental information.

### Tissue samples/Patients

CCM and arteriovenous malformation (AVM) lesions were collected from patients diagnosed clinically, neuroradiologically, and pathologically. Non-lesional controls were from patients undergoing focal epilepsy resection. All samples were collected with written informed consent under the Declaration of Helsinki and approval from the Internal Review and Ethics Committee of Beijing Tiantan Hospital.

### Mouse studies

All animal protocols were approved by the Animal Welfare and Ethics Committee of Beijing Neurosurgical Institute Laboratory. Mice were housed under specific pathogen-free conditions with a 12-hour light/dark cycle and unrestricted food and water. *Cdh5*-CreERT2; *Ccm2*^fl/fl^ (*Ccm2*^iECKO^) mice were generated by crossing *Ccm2*^fl/fl^ mice on a C57BL/6J background with *Cdh5*-CreERT2 mice. No animals or data points were excluded. Investigators were blinded during data acquisition and analysis; each animal was an experimental unit.

## RESULTS

### Study design and samples

We characterized the CCM microenvironment using 69 lesion specimens from 65 patients. Of these, 16 were profiled by single-cell RNA sequencing (scRNA-seq), 4 by Visium HD spatial transcriptomics (ST), 36 by immunohistochemistry (IHC), 8 by transmission electron microscopy (TEM) and 5 were used for primary EC immunofluorescence staining (IF) **(Figure 1A, Tables S1-S2)**. We also analyzed 16 control brain samples from patients with epilepsy (4 for scRNA-seq, 2 for ST, 6 for IHC, and 4 for primary EC IF), 15 brain arteriovenous malformation (AVM) specimens (5 for scRNA-seq, 6 for IHC, and 4 for primary EC IF), and 2 peripheral blood samples from patients with CCM. AVMs served as disease controls to distinguish CCM-specific immune features from changes shared across hemorrhage-prone cerebrovascular malformations. Integrated analysis included two external scRNA-seq datasets comprising 9 control brain samples and 1 CCM lesion ^56,57^. We validated findings in three public bulk RNA-seq cohorts comprising 25 CCM lesions and 10 control brain tissues ^58–60^.

**Figure 1.**
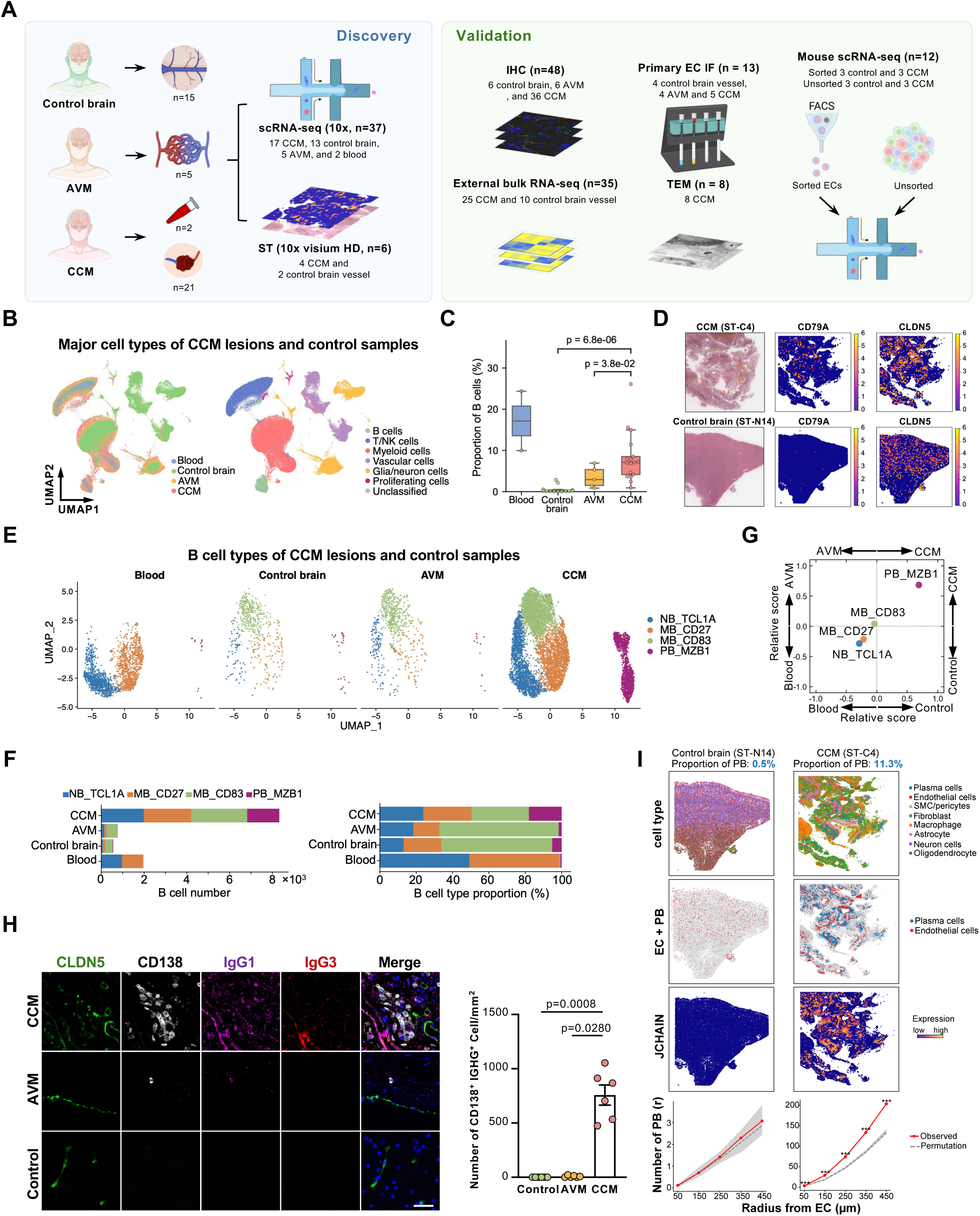
Large-scale single-cell and spatial transcriptomic profiling reveals B cell infiltration and plasma cell expansion in human CCM lesions. **(A)** Study design and sample composition. AVM, Arteriovenous Malformation. CCM, Cerebral cavernous malformations. scRNA-seq, single-cell RNA sequencing. ST, spatial transcriptomics. IHC, immunohistochemistry. TEM, transmission electron microscopy. ECs, endothelial cells. IF, immunofluorescence. **(B)** Uniform Manifold Approximation and Projection (UMAP) visualization of aggregated 300,821 single cells colored by tissue types (left) and by major cell types (right). **(C)** Box plot showing the proportion of B cells across blood, control brain, AVM, and CCM samples. **(D)** hematoxylin and eosin (H&E) staining and spatial visualization of *CD79A* (B cell marker) and *CLDN5* (EC marker) in CCM sample (ST-C4, top row) and control brain sample (ST-N14, bottom row). **(E)** UMAP plot of 11,613 B cells grouped into 4 cell subclusters, separated by tissue types. NB, naïve B cell. MB, memory B cell. PB, plasma cell. **(F)** B cell subcluster composition across tissue types. Left panel: cell counts; right panel: relative proportions of each B cell subcluster. **(G)** Two-dimensional representation of tissue-type enrichment for B cell subsets. Each quadrant corresponds to one tissue type. The position of each B cell subcluster (dot) reflects its relative proportion in that tissue type versus the others, such that a subcluster is displaced toward the tissue type in which it is enriched (Methods). Dots are color-coded by B cell subcluster. **(H)** Representative multiplexed immunohistochemistry (mIHC) imaging showing CD138^+^IgG^+^ cells, Scale bar = 50 μm. Quantification of CD138^+^IGHG^+^ cell in control, AVM and CCM. Data are presented as mean ± SEM (n = 6 per group). Three randomly selected fields were analyzed to represent each sample. Statistical analysis was performed using Kruskal-Wallis test. **(I)** Representative high-resolution Visium HD ST images of control brain and CCM showing spatial visualization of annotated cell types (row 1), distribution of EC and plasma cells (PB) (row 2), expression of the plasma cell marker *JCHAIN* (row 3). Row 4 shows EC–PB spatial proximity as the number of PB within radius r of each EC (observed, red; permutation expectation, black dashed line with gray envelope). PB were significantly enriched around EC (***p < 0.001) in CCM but not in control brain. Statistical significance was assessed using 1000 spatial permutations.

We generated whole-brain and endothelial cell-sorted scRNA-seq datasets from *Cdh5*-CreERT2; *Ccm2*^fl/fl^ mice (“*Ccm2*^iECKO^”) and wild-type littermates (n = 3 per group for each strategy; total n = 12). We integrated a published endothelial cell-sorted scRNA-seq dataset from *Ccm3*^iECKO^ and wild-type mice (n = 2 per group) ^61^.

### Transcriptional landscape reveals increased B cell infiltration and activation in human CCMs

Quality control retained 300,821 cells: 119,411 cells from CCM lesions, 15,884 from peripheral blood, 139,715 from control brain tissues, and 25,811 from AVM lesions **(Figures 1B and S1A-B)**. After batch correction ^62^, clustering identified six major cell compartments by canonical markers: vascular cells (*CLDN5*, *COL3A1*, *TAGLN*), glial/neuronal cells (*PLP1*, *MBP*, *MAP2*), myeloid cells (*P2RY12*, *CD14*, *CD68*), T/natural killer (NK) cells (*CD3D*, *NKG7*), B cells (*MS4A1*, *CD79A*), and proliferating cells (*MKI67*) **(Figures S1C-D)** ^63–68^. The automated tool Garnett supported cell-type annotations ^69^ **(Figure S1E)**.

We compared major cell-type abundance across CCM, control brain, and AVM samples **(Figures S1F-G)**. CCM and AVM lesions showed greater immune infiltration than control brains **(Figure S1F)**. B cells were more abundant in CCM lesions (mean, 8.2%; Figure 1C) than in control brains (0.1%) or AVMs (3.5%), indicating a distinctive CCM immune feature.

Differential expression analysis showed upregulated immunoglobulin genes, including *IGKC*, *IGLC2*, *IGHG1*, *IGHG3*, and *IGHG4*, and proinflammatory mediators including *S100A8* and *S100A9*, in CCM lesions compared with control brains **(Figure S1H; Table S3)**. Gene Ontology analysis showed enrichment of adaptive immune, B cell receptor, B cell-mediated, and immunoglobulin-mediated responses **(Figure S1I)**. Visium HD ST of 235,716 high-quality segmented cells from 4 CCM and 2 control samples showed broad distribution of *CD79A* B cells within CCM lesions **(Figure 1D; Table S2)**. Together, these data provide a high-resolution, multi-omic characterization of B-cell infiltration and activation in CCMs, extending previous histological observations and establishing humoral immunity as a prominent feature of the CCM immune microenvironment.

### IgG plasma cells are lesion-specific and perivascularly expanded in CCMs

To delineate the phenotypic diversity of infiltrating B cells, we reclustered the broad B cells into four transcriptionally distinct subsets: naïve B cells (NB_TCL1A), memory B cells (MB_CD27 and MB_CD83), and plasma cells (PB_MZB1) (**Figures 1E and S2A**). Plasma cells highly expressed IgG heavy-chain transcripts (*IGHG1* and *IGHG3*) and regulators of plasma cell differentiation and antibody secretion (*XBP1*, *MZB1*, and *JCHAIN*) **(Figure S2B)**. Notably, plasma cells averaged 18% of CCM B cells, exceeding levels in blood, control brain, and AVM samples **(Figures 1F and S2C)**. Tissue-enrichment projection demonstrated preferential plasma-cell accumulation in CCMs compared to other tissues **(Methods; Figure 1G)**. This finding was consistently validated across three independent bulk RNA-seq datasets, where plasma cell signature scores were repeatedly elevated in CCMs relative to controls **(Figure S2D)**. Public AVM and control-brain single-cell datasets ^70^ showed negligible plasma-cell representation **(Figure S2E)**, supporting lesion-specific expansion in CCMs.

Using the cell-potency tool CytoTRACE2 ^71^, we found that naïve B cells in CCMs were less differentiated and more transcriptionally plastic than blood-derived counterparts **(Figure S2F)**, suggesting the CCM microenvironment maintains them in an activation-ready state permissive for maturation into plasma cells. Immunoglobulin heavy-chain gene usage showed that most CCM-associated plasma cells preferentially expressed *IGHG* over *IGHA* transcripts **(Figures S2G-H)**, indicating an IgG-dominant plasma cell phenotype. Validation cohorts confirmed increased IgG-related expression in CCM lesions **(Figure S2I)**.

Multiplex immunohistochemistry (mIHC) validated CD138 IgG plasma cells, sparse in control and AVM tissues but enriched in CCM lesions. IgG signals frequently occurred near CLDN5 perivascular endothelial structures **(Figure 1H)**. Quantification confirmed higher CD138 IgG plasma-cell densities in CCM lesions **(Figure 1H)**. High-resolution Visium HD ST showed increased total and *IGHG* plasma-cell numbers along CCM vascular structures **(Figures 1I and S2J-M)**. Plasma cells, particularly *IGHG* subsets, were closer to endothelial cells in CCMs than in controls **(Figures 1I and S2N)**.

### Plasma cell abundance is associated with clinical severity in CCMs

To evaluate the impact of plasma cells on patient outcomes, we assessed plasma cell abundance across multiple modalities in CCMs. scRNA-seq analysis revealed increased plasma cell abundance in CCM lesions with subacute intracerebral hemorrhage (ICH) **(Figure S3A)**. Similarly, in spatial transcriptomic data, subacute ICH samples exhibited higher proportions of total and *IGHG* plasma cells than non-ICH CCM lesions and control brains **(Figures S3B-C)**. Bulk RNA-seq showed positive correlations between plasma cell/IgG signatures and lesion burden **(Figures S3D-E)**, while IHC in 36 CCM patients showed increased CD138 plasma cells in ICH than non-ICH CCM lesions **(Figures S3F-G)**. Plasma cell numbers within lesions also positively correlated with lesion size **(Figure S3H)**. Together, these findings establish that plasma cell accumulation is not merely a bystander feature but is tightly linked to CCM severity, hemorrhagic risk, and tissue remodeling—highlighting plasma cells as clinically relevant biomarkers and potential therapeutic targets.

### Plasma cells are linked to endothelial senescence in human CCM lesions

Endothelial senescence contributes to vascular dysfunction, but its immune-related regulation in human CCMs remains incompletely defined ^1–10,72,73^. Recent studies report that IgG accumulates in aging tissues and promotes macrophage and microglial senescence ^11^, which is associated with the activated NF-κB pathway. However, whether IgG can induce endothelial senescence has not been explored.

To evaluate endothelial senescence in human CCMs, we first reclustered vascular cells into arterial (Art_VEGFC), capillary (Cap_MFSD2A), venous (Vn_ACKR1), pericyte (PC_KCNJ8), smooth muscle (SMC_ACTA2), and fibroblast (FB_DCN and FB_IGFBP5) populations **(Figures 2A and S4A-B)**. Across endothelial subtypes, CCM samples showed significantly elevated expression of multiple established senescence-associated gene programs, including senescence-associated secretory phenotype (SASP) signatures (EC_SASP, SenMayo, and Reactome R-HSA-2559582) and the Gene Ontology cellular senescence term (GO:0090398), compared with control brains ^74–77^ **(Figures 2B and S4C; Table S4)**. CCM endothelial cells upregulated SASP cytokines and chemokines, including *IL6*, *CCL2*, *CCL3*, *CCL5*, *TGFB1*, *IGFBP2*, *CXCL2*, and *ICAM1* **(Figures 2C and S5)**.

**Figure 2.**
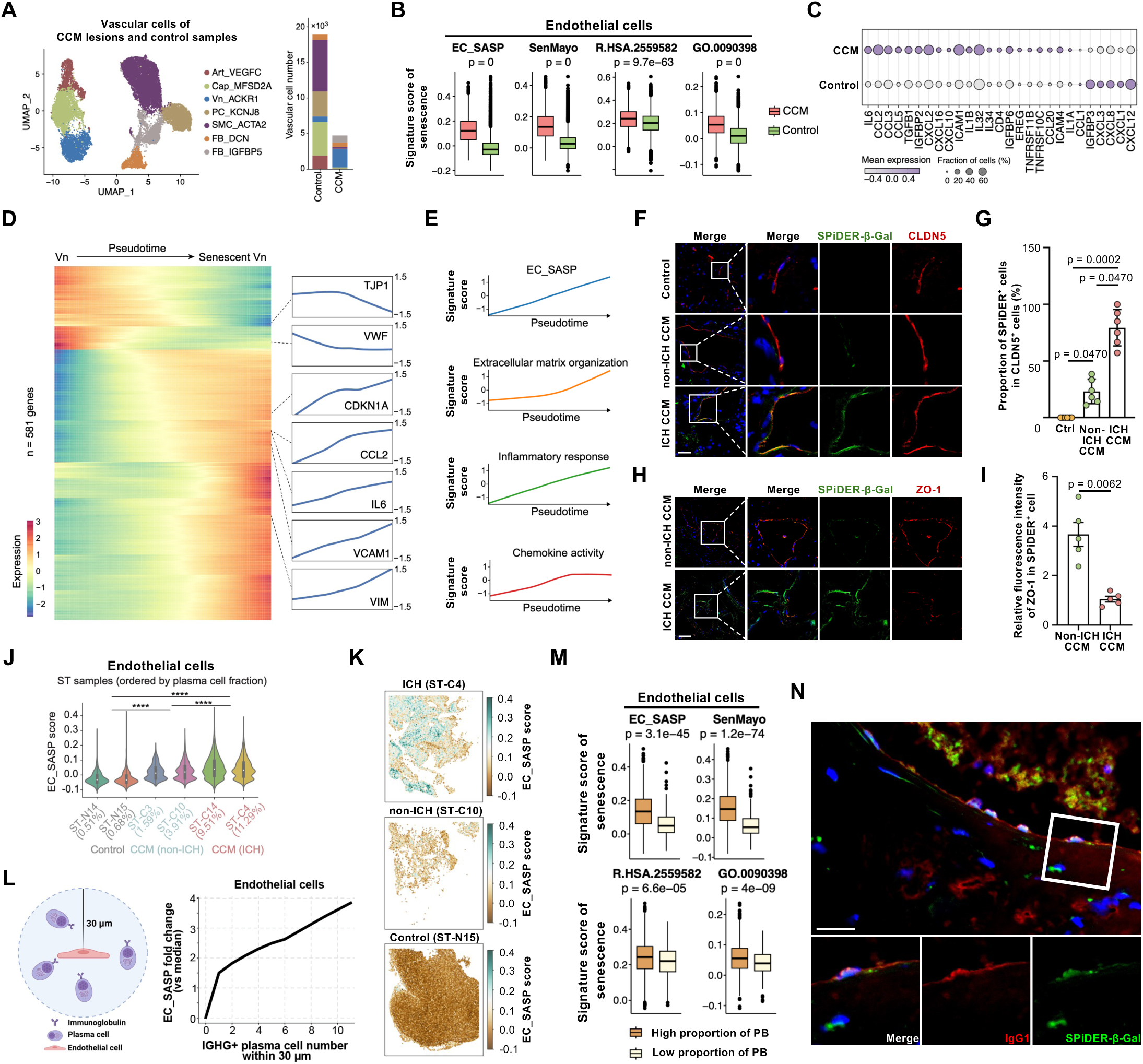
Endothelial cells display elevated senescence phenotypes in CCM lesions. **(A)** UMAP plot of 26,780 vascular cells grouped into seven cell subclusters, separated by tissue types (control brains and CCM lesions, left). Number of vascular subclusters (right). **(B)** Box plots showing the distribution of senescence-associated signature scores, including EC_SASP (De Cecco et al.), SenMayo, Reactome senescence pathway (R.HSA.2559582), and Gene Ontology (GO) senescence signature (GO.0090398), in ECs from CCM lesions compared with control brain samples. SASP: senescence-associated secretory phenotype. **(C)** Dot plot showing the expression of inflammatory chemokines and cytokines in ECs from CCM lesions compared to control brains. **(D)** Pseudotime analysis during the transition from venous ECs to senescent venous ECs, showing DEGs (left), and representative gene expression patterns (right). **(E)** Regression curves showing the trajectory of gene signature scores along pseudotime. **(F)** Representative images of control, subacute ICH CCM, and non-ICH CCM samples stained with SPiDER-β-Gal and CLDN5. Scale bar = 50 μm. **(G)** Quantification of the proportion of SPiDER-β-Gal-positive cells among CLDN5-positive endothelial cells. Data are presented as mean ± SEM (n = 6 per group). Three randomly selected fields were analyzed to represent each sample. Statistical analyses were performed using one-way ANOVA. **(H)** Representative images of subacute ICH CCM and non-ICH CCM samples stained with SPiDER-β-Gal and ZO-1. Scale bar = 50 μm. **(I)** Quantification of relative fluorescence intensity of ZO-1. Data are presented as mean ± SEM (n = 5 per group). Three randomly selected fields were analyzed to represent each sample. Statistical analysis was performed using unpaired t-test. **(J)** Violin plots showing EC_SASP scores in ECs across ST samples ordered by PB cell fraction. Wilcoxon rank-sum test. **** p < 0.0001. **(K)** Spatial expression of EC_SASP scores across ICH CCM (ST-C4), non-ICH CCM (ST-C10), and control brain (ST-N15) samples. **(L)** Schematic (left) and line plot (right) illustrating the relationship between EC senescence and local IGHG plasma cell density. For each EC, IGHG plasma cells were counted within a 30-μm radius (schematic), and the per-cell EC_SASP score was normalized to the median EC_SASP score across all ECs (Methods). The LOWESS-smoothed curve shows that EC_SASP score increases with the number of neighboring IGHG plasma cells. **(M)** Box plots showing the distribution of senescence-associated signature scores in CCM lesions stratified by PB abundance. Samples were categorized into high-PB and low-PB groups based on the median proportion of PBs among all cells. **(N)** Representative images showing the colocalization of IgG1 and SPiDER-β-Gal staining. Scale bar: 50 μm.

We constructed a pseudotime trajectory ^78^ from venous to senescence-associated venous endothelial states and identified 581 dynamically regulated genes **(Figure 2D; Table S5)**. Along this trajectory, SASP cytokines (*IL6* and *CCL2*), the cell-cycle inhibitor *CDKN1A*, the adhesion molecule *VCAM1*, and the mesenchymal marker *VIM* progressively increased, whereas the tight-junction component *TJP1* and the venous identity gene *VWF* decreased **(Figures 2D-E)**. These data link senescence-associated endothelial states with inflammatory activation, mesenchymal-like features, and reduced vascular integrity markers.

Histological validation using SPiDER-β-galactosidase (SPiDER-β-Gal) staining confirmed increased endothelial senescence in CCM lesions, with higher levels in subacute ICH lesions than in non-ICH lesions **(Figures 2F-G)**. Tight-junction protein ZO-1 expression was markedly reduced in ICH samples **(Figures 2H-I**), suggesting impaired barrier function. Spatial Visium HD analysis further supported these findings, showing elevated endothelial senescence scores in CCM-ICH samples **(Figures 2J-K and S5)**. As described above, ICH lesions also exhibited higher plasma cell abundance **(Figures S3A-C)**. For each endothelial cell, we quantified the number of *IGHG*^+^ plasma cells within a 30-μm radius and correlated this measure to its senescence score **(Figure 2L)**. Endothelial cells located near higher numbers of *IGHG*^+^ plasma cells exhibited higher senescence scores. Consistently, endothelial cells from plasma cell-rich CCM lesions showed higher senescence signatures and increased inflammatory chemokine expression, with the strongest effects observed in venous endothelial cells **(Figures 2M and S4D-E)**. Co-staining for IgG1 and SPiDER-β-Gal further demonstrated spatial overlap between IgG deposition and senescent cells **(Figure 2N)**. Together, these findings identify endothelial senescence as a feature of human CCMs spatially associated with local plasma cell accumulation and hemorrhage-associated phenotypes.

### IgG induces endothelial senescence through an mTOR-dependent NF-**κ**B pathway

To determine whether IgG can directly promote endothelial senescence, we treated human umbilical vein endothelial cells (HUVECs) with purified native IgG using a concentration established in a recent study ^11^. IgG exposure markedly increased senescence-associated β-galactosidase (SA-β-Gal) activity and P21 expression **(Figures 3A-D)**, together with upregulation of canonical SASP factors **(Figure 3E)**. Because endothelial senescence is associated with impaired junctional integrity ^7^, we next examined ZO-1 expression after IgG treatment. IgG reduced ZO-1 abundance and disrupted junctional organization **(Figures 3F-G)**, indicating compromised endothelial barrier architecture. These findings show that IgG induces endothelial senescence and tight-junction deterioration in vitro, recapitulating features of human CCM lesions.

**Figure 3.**
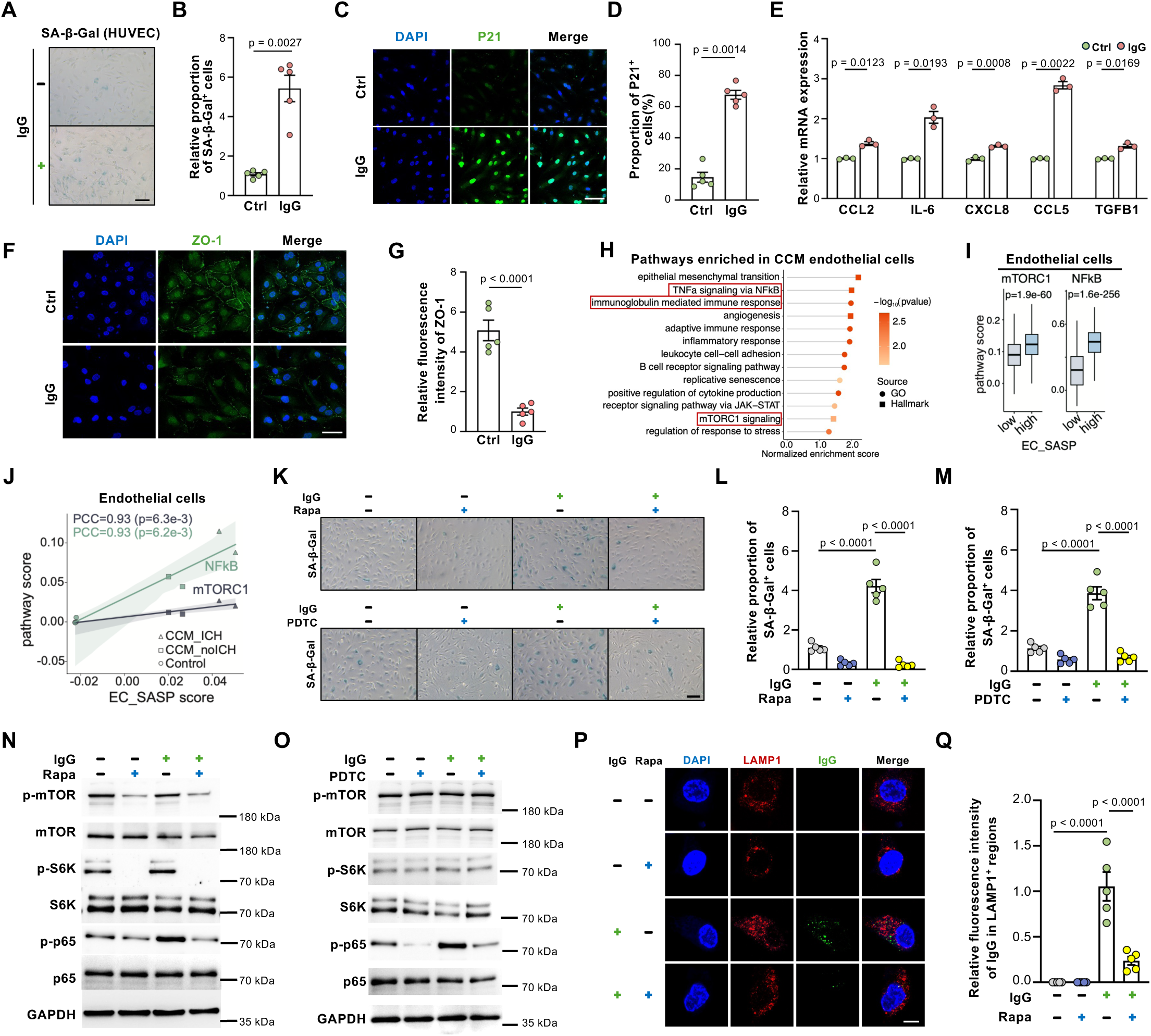
IgG induces endothelial cell senescence through mTOR-dependent activation of the NF-κB pathway. **(A)** Representative SA-β-Gal staining in HUVECs treated with vehicle control (Ctrl) or IgG. Scale bar = 50 μm. **(B)** Quantification of the proportion of SA-β-Gal positive cells in IgG treatment and Ctrl group. Data are presented as mean ± SEM (n = 5 biological replicates). Three randomly selected fields were analyzed to represent biological replicate. Statistical analysis was performed using unpaired t-test. **(C)** Immunofluorescence staining of P21 in HUVECs treated with Ctrl or IgG. Scale bar = 50 μm. **(D)** Quantification of the proportion of P21-positive cells in IgG treatment and Ctrl group. Data are presented as mean ± SEM (n = 5 biological replicates). Three randomly selected fields were analyzed to represent each biological replicate. Statistical analysis was performed using unpaired t-test. **(E)** Quantitative reverse transcription PCR (RT-PCR) analysis of *CCL2*, *IL6*, *CXCL8*, *CCL5*, and *TGFB1* in HUVECs treated with IgG or Ctrl. Data are presented as mean ± SEM (n = 3 biological replicates). Statistical analysis was performed using unpaired t-test. **(F)** Immunofluorescence staining of ZO-1 in HUVECs treated with Ctrl or IgG. Scale bar = 50 μm. **(G)** Quantification of the fluorescence intensity of ZO-1. Data are presented as mean ± SEM (n = 5 biological replicates). Three randomly selected fields were analyzed to represent each biological replicate. Statistical analysis was performed using unpaired t-test. **(H)** Pathway enrichment analysis of genes upregulated in ECs from CCM lesions compared with control brains, based on scRNA-seq data. Terms are colored by enrichment significance (−log P value) and shaped by gene-set source (GO or Hallmark), and ranked by normalized enrichment score. **(I)** Boxplots displaying mTORC1 and NF-κB pathway activity in ECs stratified by EC_SASP scores in CCM scRNA-seq data. Wilcoxon rank-sum test. **(J)** Correlation between EC_SASP scores and mTORC1 or NF-κB pathway scores across ST samples. Each point represents one ST sample. Lines show linear regression fits (PCC, Pearson correlation coefficient). Both pathways were positively correlated with EC_SASP score. **(K)** Representative SA-β-Gal staining images showing the effect of IgG, rapamycin (top), and PDTC (bottom) on endothelial senescence. Scale bar = 50 μm. **(L-M)** Quantification of the proportion of SA-β-Gal-positive cells in response to IgG treatment, with and without Rapamycin (L) or PDTC (M) treatment. Data are presented as mean ± SEM (n = 5 biological replicates). Three randomly selected fields were analyzed to represent each biological replicate. Statistical analysis was performed using one-way ANOVA. Relative to IgG^-^/rapamycin^-^ or IgG^-^/PDTC^-^ group. **(N-O)** Western blot showing the expression levels of mTOR, p-mTOR, S6K, p-S6K, p65, and p-p65 in HUVECs in response to IgG treatment, with and without rapamycin (N) or PDTC (O) treatment. GAPDH is shown as a loading control. **(P)** Immunofluorescence staining of IgG and LAMP1 in HUVECs treated with or without IgG or Rapamycin. Scale bar = 10 μm. **(Q)** Quantification of fluorescence intensity of IgG in LAMP1-positive area among groups. Data are presented as mean ± SEM (n = 5 biological replicates). Three randomly selected fields were analyzed to represent each biological replicate. Statistical analysis was performed using one-way ANOVA. Relative to IgG^+^/rapamycin^-^ group.

We next examined mechanisms underlying IgG-induced endothelial senescence. Gene set enrichment analysis of endothelial cells from CCM lesions revealed enrichment of senescence-associated programs together with activation of mTORC1 and NF-κB signaling compared with control brains **(Figure 3H),** consistent with previous reports of mTOR activation in CCM ^79,80^, whereas NF-κB is a well-established mediator of inflammatory senescence ^81^. To determine whether these pathways were linked to endothelial senescence, we compared endothelial cells with high and low EC_SASP scores. Endothelial cells with elevated EC_SASP scores exhibited significantly greater mTORC1 and NF-κB pathway activity**(Figure 3I)**, and spatial transcriptomic analysis independently confirmed positive correlations between EC_SASP scores and both pathways across CCM lesions **(Figure 3J)**.

Because IgG has been linked to NF-κB–dependent senescence-like programs in myeloid cells ^11^, and mTORC1 regulates lysosomal catabolism ^18,82^, we investigated whether IgG-induced endothelial senescence depends on an mTOR-lysosome-NF-κB signaling axis. We first examined the contribution of mTOR and NF-κB signaling to IgG-induced endothelial senescence using pharmacological inhibitors. HUVECs were treated with IgG in the presence or absence of inhibitors targeting mTOR or NF-κB. Inhibition of either mTOR or NF-κB significantly attenuated IgG-induced endothelial senescence **(Figures 3K-M)**, indicating that both pathways are required for the senescence response. We next determined the signaling relationship between mTOR and NF-κB. IgG treatment increased phosphorylation of the NF-κB subunit p65 but did not alter phosphorylation of mTOR or its downstream target ribosomal protein S6 kinase (S6K) **(Figures 3N-O and S6A-B)**, suggesting that IgG directly activates NF-κB without further stimulating mTOR signaling. However, mTOR inhibition with rapamycin markedly attenuated IgG-induced p65 phosphorylation **(Figures 3N-O and S6A-B)**, suggesting mTOR is upstream of NF-κB activation.

Because mTOR is a central regulator of lysosomal function, we next asked whether it controls intracellular IgG handling. Immunofluorescence analysis demonstrated that rapamycin reduced intracellular IgG accumulation within lysosome-associated membrane protein 1 (LAMP1) ^+^ lysosomal compartments **(Figures 3P-Q)**, whereas NF-κB inhibition had no effect on intracellular IgG levels **(Figures S6C-D)**, indicating that lysosomal mTOR regulates IgG handling upstream of NF-κB activation.

Finally, to determine whether impaired lysosomal degradation is sufficient to promote endothelial senescence, we inhibited lysosomal acidification with bafilomycin A1 (BafA1). BafA1 treatment increased intracellular IgG accumulation, NF-κB activation, and endothelial senescence, all of which were attenuated by rapamycin **(Figures S7A-D)**. Together, these findings support a model in which mTOR-dependent lysosomal dysfunction promotes intracellular IgG accumulation, leading to NF-κB activation and endothelial senescence.

### CCM loss-induced mTOR activation sensitizes endothelial cells to accumulate lysosomal IgG promoting endothelial senescence

Previous studies have shown elevated mTOR activity after *CCM1*/*2*/*3* loss-of-function or *PIK3CA* gain-of-function mutations in CCM lesions ^83^. We hypothesized that sustained mTOR activation in CCM endothelium enhances susceptibility to IgG-associated senescence by altering intracellular IgG handling.

To test this hypothesis, we depleted *CCM2* in HUVECs and examined the effects of IgG exposure in the presence or absence of rapamycin. *CCM2* knockdown increased phosphorylation of mTOR and its downstream target S6K regardless of IgG treatment **(Figures 4A and S8A)**, confirming constitutive mTOR activation. Although *CCM2* knockdown alone modestly increased endothelial senescence, an effect attenuated by rapamycin, suggesting that elevated mTOR activity contributes to basal endothelial senescence **(Figures 4B-C)**, IgG exposure markedly enhanced intracellular IgG accumulation, NF-κB activation (p65 phosphorylation), and SA-β-Gal activity in *CCM2*-deficient cells, all of which were attenuated by mTOR inhibition **(Figures 4A-D and S8A**), indicating that mTOR activation amplifies IgG-induced endothelial dysfunction.

**Figure 4.**
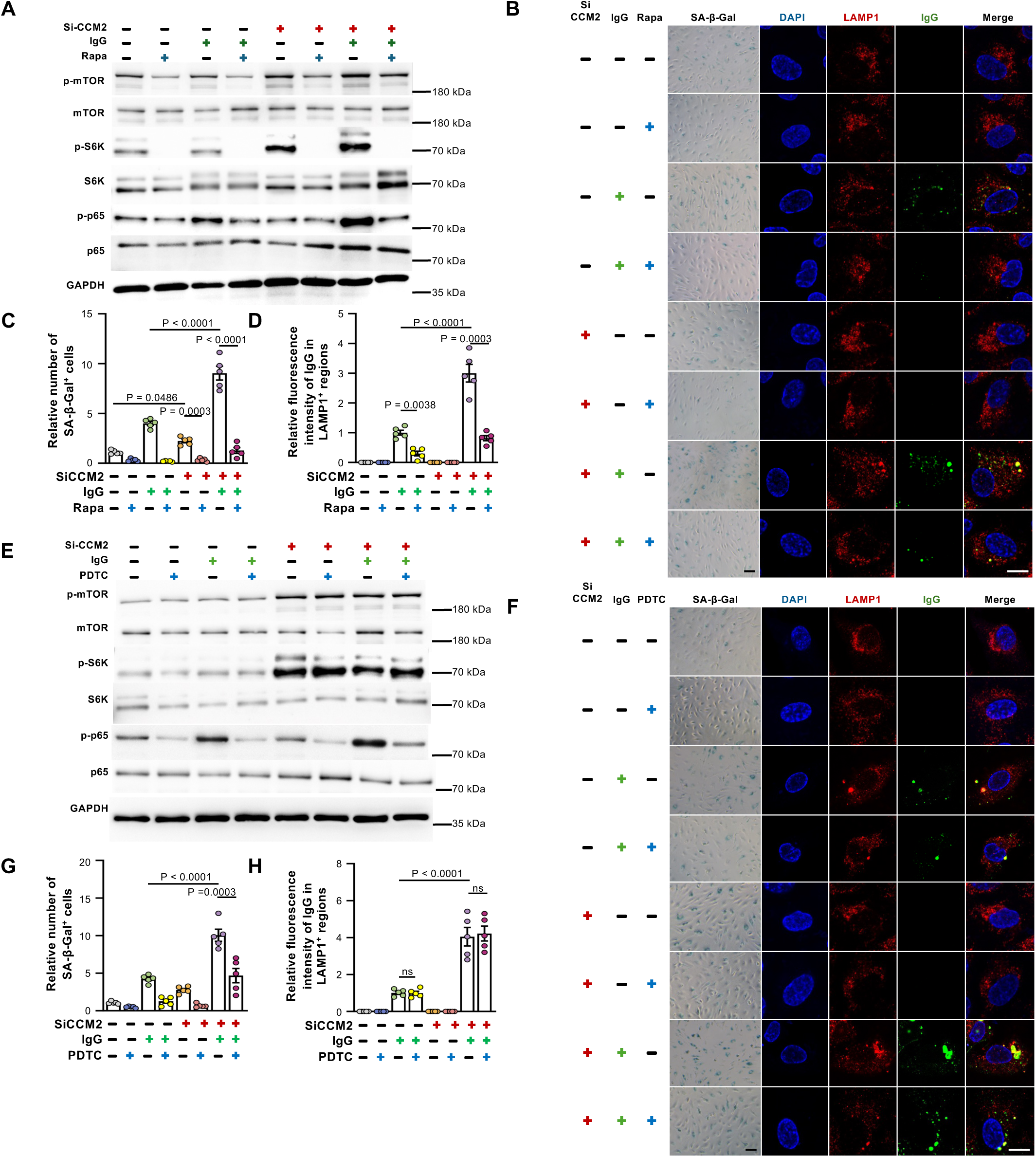
mTOR activation in *CCM2*-deficient endothelial cells converges with IgG stimulation accelerating endothelial cell senescence. **(A)** Representative blots of HUVECs under eight experimental conditions. The same groups generated by the combination of IgG (−/+), rapamycin (Rapa) (−/+), and *CCM2* knockdown (−/+) were subjected to Western blotting (WB) to assess protein expression levels of mTOR, p-mTOR, S6K, p-S6K, p65, and p-p65. **(B)** HUVECs were treated under eight experimental conditions generated by the combination of three factors: IgG (−/+), rapamycin (−/+), and *CCM2* knockdown (−/+). SA-β-Gal staining and immunofluorescence staining for IgG and LAMP1 were performed to assess endothelial senescence and IgG accumulation, respectively. Scale bar (SA-β-Gal) = 50 μm. Scale bar (IF) = 10 μm. **(C)** Quantification of the number of SA-β-Gal positive cells. Data are presented as mean ± SEM (n = 5 biological replicates). Three randomly selected fields were analyzed to represent each sample. Statistical analysis was performed using one-way ANOVA. Relative to siCCM2^-^/IgG^-^/Rapamycin^-^ group. **(D)** Quantification of fluorescence intensity of IgG in LAMP1-positive area among groups. Data are presented as mean ± SEM (n = 5 biological replicates). Three randomly selected fields were analyzed to represent each sample. Statistical analysis was performed using one-way ANOVA. Relative to siCCM2^-^/IgG^+^/Rapamycin^-^ group. **(E)** Representative blots of HUVECs under eight experimental conditions. The same groups generated by the combination of IgG (−/+), PDTC (−/+), and *CCM2* knockdown (−/+) were subjected to WB to assess protein expression levels of mTOR, p-mTOR, S6K, p-S6K, p65, and p-p65. **(F)** HUVECs were treated under eight experimental conditions generated by the combination of three factors: IgG (−/+), PDTC (−/+), and *CCM2* knockdown (−/+). SA-β-Gal staining and immunofluorescence staining for IgG and LAMP1 were performed to assess endothelial senescence and IgG accumulation, respectively. Scale bar (SA-β-Gal) = 50 μm. Scale bar (IF) = 10 μm. **(G)** Quantification of the percentage of SA-β-Gal positive cells. Data are presented as mean ± SEM (n = 5 biological replicates). Three randomly selected fields were analyzed to represent each sample. Statistical analysis was performed using one-way ANOVA. Relative to siCCM2^-^/IgG^-^/PDTC^-^ group. **(H)** Quantification of fluorescence intensity of IgG in LAMP1-positive area among groups. Data are presented as mean ± SEM (n = 5 biological replicates). Three randomly selected fields were analyzed to represent each sample. Statistical analysis was performed using one-way ANOVA. Relative to siCCM2^-^/IgG^+^/PDTC^-^ group.

To determine whether NF-κB mediates this response downstream of mTOR, we inhibited NF-κB during IgG treatment. NF-κB inhibition significantly attenuated endothelial senescence in *CCM2*-deficient cells **(Figures 4E-H and S8B)**, supporting that IgG-associated senescence remains NF-κB dependent even in the setting of elevated mTOR activity. Similar results were obtained in human brain microvascular endothelial cells (HBMECs), confirming that this mechanism is not restricted to HUVECs **(Figures S9A-D)**.

We further investigated whether impaired lysosomal degradation contributes to lysosomal IgG accumulation in *CCM2*-deficient cells. Immunofluorescence analysis demonstrated a marked increase in intracellular IgG accumulation in CCM2 LOF HUVECs compared with control **(Figures S10A-D)**. Thus, CCM2 LOF HUVECs exhibited concomitant accumulation of intracellular IgG. Inhibition of lysosomal acidification with BafA1 further increased lysosomal IgG accumulation, whereas rapamycin reduced lysosomal IgG levels **(Figures S10A-D)**. These findings suggest that mTOR-mediated lysosomal regulation contributes to altered IgG handling in CCM loss-of-function endothelium.

Collectively, these results support a model in which CCM loss-of-function establishes a constitutively activated mTOR state that impairs lysosomal IgG processing, rendering endothelial cells hypersensitive to extrinsic IgG. Persistent intracellular IgG then promotes NF-κB-dependent endothelial senescence, providing a mechanistic link between CCM genetic defects and humoral immune–mediated vascular dysfunction.

### CCM loss-associated mTOR activation impaired lysosomal acifidication and IgG processing in HUVECs and CCM-derived endothelial cells promoting intracellular IgG accumulation

Mammalian cells can acquire exogenous amino acids through endocytosis and lysosomal catabolism of extracellular proteins ^18^. mTORC1 suppresses the nutritional use of extracellular proteins by directly controlling the catabolic activity of lysosomes ^18^. Detailedly, mTORC1 inactivation triggers assembly of the V-ATPase at lysosomal membranes, which acidifies the organelle lumen and activates lysosomal proteases ^18^. By increasing catabolic activity throughout the lysosomal population, cells initiate degradation of protein contents from extra-and intracellular sources that were accumulated in lysosomes; mTORC1 activation blocks lysosomal degradation of extracellular proteins by suppressing V-ATPase-mediated acidification of lysosomes assembly ^18^. A previous report indicated that defective autophagy is a key feature of cerebral cavernous malformations ^84^. Our preceding experiments implicated mTOR-dependent lysosomal regulation in endothelial IgG accumulation, we next directly examined lysosomal acidification and IgG deposition in CCM2 LOF HUVECs and CCM-derived endothelial cells.

Firstly, to determine whether CCM loss of function affects lysosomal acidification in endothelial cells, we measured lysosomal pH in HUVECs using the ratiometric fluorescent probe LysoSensor Yellow/Blue DND-160. Following in situ calibration of the fluorescence ratio against buffers of defined pH, absolute lysosomal pH values were calculated for each experimental condition. Compared with control HUVECs, CCM-deficient cells exhibited a significant increase in lysosomal pH, indicating impaired lysosomal acidification and lysosomal alkalinization **(Figures 5A-B and S10E)**. Notably, treatment with rapamycin significantly reduced the elevated lysosomal pH in CCM-deficient HUVECs, indicating a partial restoration of lysosomal acidification. In contrast, bafilomycin A1 treatment further increased lysosomal pH in CCM-deficient cells, consistent with inhibition of vacuolar H -ATPase–dependent lysosomal acidification. These findings demonstrate that CCM loss of function disrupts lysosomal acidification in endothelial cells and suggest that modulation of the autophagy–lysosomal pathway can influence this lysosomal pH defect **(Figures 5A-B)**.

**Figure 5.**
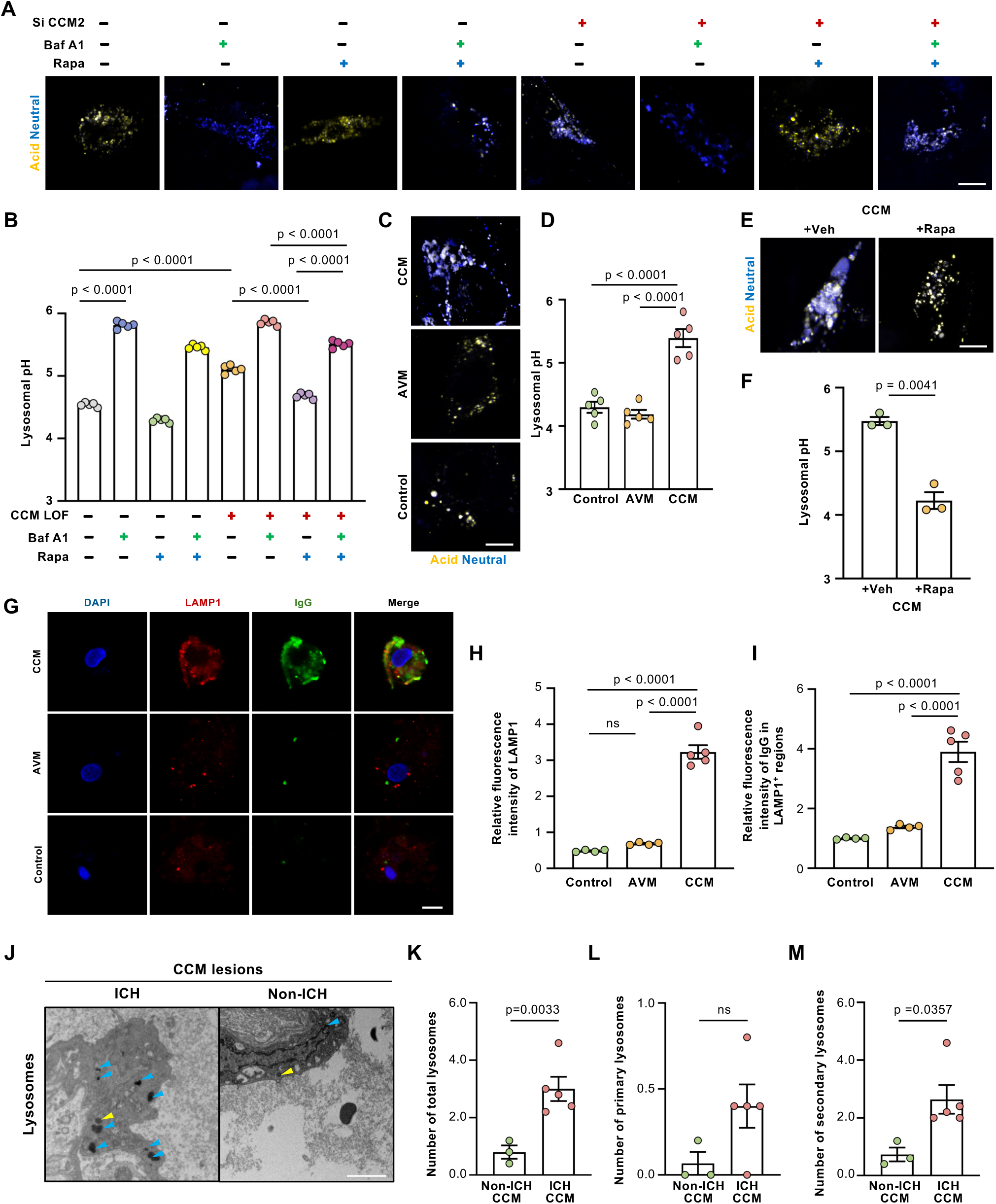
CCM loss-associated mTOR activation impairs lysosomal acidification and IgG processing in endothelial cells. **(A)** Representative ratiometric lysosomal pH images of endothelial cells under eight experimental conditions generated by the combination of CCM loss of function (CCM LOF), bafilomycin A1 (Baf A1), and rapamycin (Rapa). Yellow and blue signals represent relatively acidic and less acidic/neutral lysosomal compartments, respectively. **(B)** Quantification of lysosomal pH under the eight experimental conditions shown in (A). Data are presented as mean ± SEM. Statistical analysis was performed using one-way ANOVA with comparisons as indicated. Data are presented as mean ± SEM (n = 5 cells). Each data point represents one individual cell. **(C)** Representative ratiometric lysosomal pH images of primary endothelial cells isolated from cerebral cavernous malformations (CCM), brain arteriovenous malformations (AVM), and non-lesional brain tissues obtained during epilepsy surgery (Control). **(D)** Quantification of lysosomal pH in primary endothelial cells from Control, AVM, and CCM samples. Data are presented as mean ± SEM. Statistical analysis was performed using one-way ANOVA. Data are presented as mean ± SEM (n = 5 cells). Each data point represents one individual cell. **(E)** Representative ratiometric lysosomal pH images of primary CCM-derived endothelial cells treated ex vivo with vehicle (Veh) or rapamycin (Rapa). **(F)** Quantification of lysosomal pH in vehicle-and rapamycin-treated CCM-derived endothelial cells. Data are presented as mean ± SEM. Statistical analysis was performed using an unpaired t-test. Data are presented as mean ± SEM (n = 5 cells). Each data point represents one individual cell. **(G)** Representative immunofluorescence images of IgG and the lysosomal marker LAMP1 in primary endothelial cells isolated from CCM lesions, AVMs, and non-lesional control brain tissues. Nuclei were counterstained with DAPI. Scale bar = 10 μm. **(H)** Quantification of relative LAMP1 fluorescence intensity in primary endothelial cells from Control, AVM, and CCM samples. Data are presented as mean ± SEM. Statistical analysis was performed using one-way ANOVA. Control, n = 4 independent donors; AVM, n = 4; CCM, n = 5. Each point represents one independent donor. **(I)** Quantification of IgG fluorescence intensity within LAMP1-positive regions in primary endothelial cells from Control, AVM, and CCM samples. Data are presented as mean ± SEM. Statistical analysis was performed using one-way ANOVA. Control, n = 4 independent donors; AVM, n = 4; CCM, n = 5. Each point represents one independent donor. **(J)** Representative transmission electron microscopy (TEM) images of primary lysosome (yellow arrowhead) and secondary lysosomes (blue arrowheads) in endothelial cells. Scale bars: 2 μm. **(K-M)** Quantitative analysis of total lysosomes, primary and secondary lysosomes between ICH CCM and non-ICH CCM groups. Data are presented as mean ± SEM (n = 3 in non-ICH CCM group, n = 5 in ICH CCM group). Five randomly selected fields were analyzed to represent each sample. Statistical analysis for the total lysosomes was performed using unpaired t-test, statistical analysis for the primary and secondary lysosomes was performed using Wilcoxon rank-sum test.

Secondly, to determine whether the lysosomal acidification defect observed following CCM loss of function in vitro is recapitulated in human lesions, we isolated endothelial cells from surgically resected human CCM specimens and assessed lysosomal pH using the ratiometric fluorescent probe LysoSensor Yellow/Blue DND-160. Endothelial cells isolated from surgically resected brain AVMs and non-lesional normal cerebral vessels were analyzed in parallel as disease and physiological controls, respectively. Following in situ calibration of the LysoSensor fluorescence ratio, absolute lysosomal pH was determined in the isolated endothelial cells. Compared with endothelial cells from normal cerebral vessels, CCM-derived endothelial cells exhibited a significant increase in lysosomal pH, indicating impaired lysosomal acidification and lysosomal alkalinization. In contrast, endothelial cells derived from AVM specimens did not show a comparable increase in lysosomal pH. These findings demonstrate that defective lysosomal acidification is present in endothelial cells from human CCM lesions and suggest that this phenotype is relatively specific to CCM pathology rather than a general feature of cerebral vascular malformations **(Figures 5C-D)**.

To further determine whether the lysosomal acidification defect in human CCM endothelium is therapeutically reversible, endothelial cells isolated from surgically resected human CCM lesions were treated with rapamycin and analyzed using LysoSensor Yellow/Blue DND-160. Absolute lysosomal pH was quantified following in situ calibration of the fluorescence ratio. Consistent with our observations in CCM-deficient HUVECs, endothelial cells derived from human CCM lesions exhibited elevated lysosomal pH, indicating impaired lysosomal acidification. Notably, rapamycin treatment significantly reduced lysosomal pH in CCM-derived endothelial cells, partially restoring lysosomal acidification toward the level observed in control endothelial cells. These findings suggest that the lysosomal acidification defect associated with CCM pathology is at least partially reversible by mTOR inhibition**(Figures 5E-F)**.

Finally, to further validate our findings in human disease, we isolated patient-derived primary endothelial cells from freshly resected CCM lesions, AVMs, and non-lesional brain tissues obtained during epilepsy surgery to investigate intracellular IgG accumulation and lysosomal alterations. Immunofluorescence staining revealed substantially increased intracellular IgG accumulation in CCM-derived endothelial cells compared with endothelial cells derived from AVMs or non-lesional epilepsy controls **(Figures 5G-I)**. Notably, CCM-derived endothelial cells also exhibited increased LAMP1 fluorescence, indicative of an expanded or remodeled LAMP1-positive lysosomal compartment **(Figures 5G-I)**. Thus, the increased intracellular IgG accumulation in CCM endothelium was accompanied by an increase in the abundance of LAMP1-positive lysosomes. These findings further validate our observations in CCM2-deficient cultured endothelial cells and demonstrate that IgG accumulation and lysosomal alterations are recapitulated in endothelial cells derived from human CCM lesions.

In addition, we performed transmission electron microscopy (TEM) of CCM lesions to examine whether lysosomal remodeling was observed in human CCM lesions. Secondary lysosomes is a recognized ultrastructural feature of altered lysosomal clearance and senescence-associated lysosomal expansion ^85^. Compared with non-ICH CCM lesions, ICH-associated CCM lesions showed a marked increase in total and secondary lysosomes within endothelial cells, whereas primary lysosome abundance was not significantly different between groups **(Figures 5J-M)**. These ultrastructural findings are consistent with impaired lysosomal processing in hemorrhagic CCM lesions and support our mechanistic model that mTOR-dependent lysosomal dysfunction promotes intracellular IgG accumulation and endothelial senescence.

Together, these findings demonstrate that CCM loss is associated with defective lysosomal acidification that can be restored by mTOR inhibition. This lysosomal dysfunction is accompanied by increased lysosomal abundance, intracellular IgG accumulation, and secondary lysosomes expansion in human CCM endothelium, providing a mechanistic link between CCM loss-associated mTOR activation and impaired endothelial IgG processing.

### Plasma cell accumulation and endothelial senescence in CCM mouse models

To determine whether the plasma cell enrichment and endothelial senescence observed in human CCMs are conserved in vivo, we analyzed inducible endothelial-specific *Ccm2* knockout *Ccm2*^iECKO^ mice. Whole-brain scRNA-seq identified nine major cell populations **(Figures S11A)**. *Ccm2*^iECKO^ mice showed increased B cell abundance compared with wild-type (WT) controls **(Figure S11B)**, accompanied by elevated expression of plasma cell marker genes (*Sdc1* and *Mzb1*) and IgG heavy-chain genes (*Ighg1*, *Ighg2b*, and *Ighg3*) **(Figure S11C)**, consistent with plasma cell enrichment.

Because endothelial cells were relatively underrepresented in the whole-brain scRNA-seq dataset, we enriched them by fluorescence-activated cell sorting (FACS) before scRNA-seq. Clustering of sorted endothelial cells (*Ccm2*^iECKO^: 9,865 cells; WT: 7,926 cells) resolved distinct endothelial subtypes, including arterial, capillary-arterial, capillary-venous, venous, and *Ccnd1* populations **(Figures S11D-F)**. Across these populations, endothelial cells from *Ccm2*^iECKO^ mice showed upregulation of senescence-associated transcriptional programs **(Figure S11G)**, closely mirroring the human CCM findings. This signature was broadly distributed across endothelial subtypes **(Figure S11H)**, indicating a widespread shift toward senescence-associated endothelial states in the CCM vasculature. Endothelial cells from *Ccm2*^iECKO^ mice also showed increased mTORC1 and NF-κB pathway activity relative to wild-type controls **(Figure S11I)**, particularly among cells with high EC_SASP scores **(Figure S11J)**, suggesting conservation across species.

We further analyzed an independent scRNA-seq dataset from a *Ccm3*^iECKO^ mouse model ^61^. Consistent with our *Ccm2*^iECKO^ data, senescence-associated gene signatures were increased in endothelial cells from *Ccm3*^iECKO^ mice compared with wild-type controls **(Figure S11K)**, supporting the conservation of endothelial senescence across genetically distinct CCM models.

At the tissue level, IHC analysis showed increased P21 expression and IgG deposition in endothelial regions of *Ccm2*^iECKO^ mice compared with WT controls **(Figures S11L-N)**. SPiDER-β-Gal staining further demonstrated increased endothelial senescence in *Ccm2*^iECKO^ mice **(Figures S11O-P)**. Senescent endothelial cells in CCM mice showed markedly reduced ZO-1 expression, indicating compromised tight-junction integrity **(Figures S11Q-R)**. Together, these data demonstrate that plasma cell enrichment in murine CCM models coincides with widespread endothelial senescence and tight-junction disruption, supporting a conserved association between plasma cell/IgG enrichment, endothelial senescence, and vascular barrier impairment in CCM models.

### Rapamycin suppressed lesion progression with reduced endothelial IgG accumulation and marked attenuation of endothelial senescence

To test whether mTORC1 signaling regulates endothelial IgG accumulation and senescence in vivo, we treated adult *Ccm2*^iECKO^ mice with magnetic resonance imaging (MRI)-confirmed CCM lesions with mTOR inhibitor rapamycin or vehicle with repeat MRI assessment after one month **(Figure 6A)**. Consistent with prior studies ^83^, rapamycin suppressed lesion progression **(Figures 6B-D)**. Histological analyses confirmed inhibition of mTOR signaling and showed reduced endothelial IgG accumulation together with marked attenuation of endothelial senescence **(Figures 6E-I)**, supporting a role for mTOR in regulating endothelial IgG handling in vivo.

**Figure 6.**
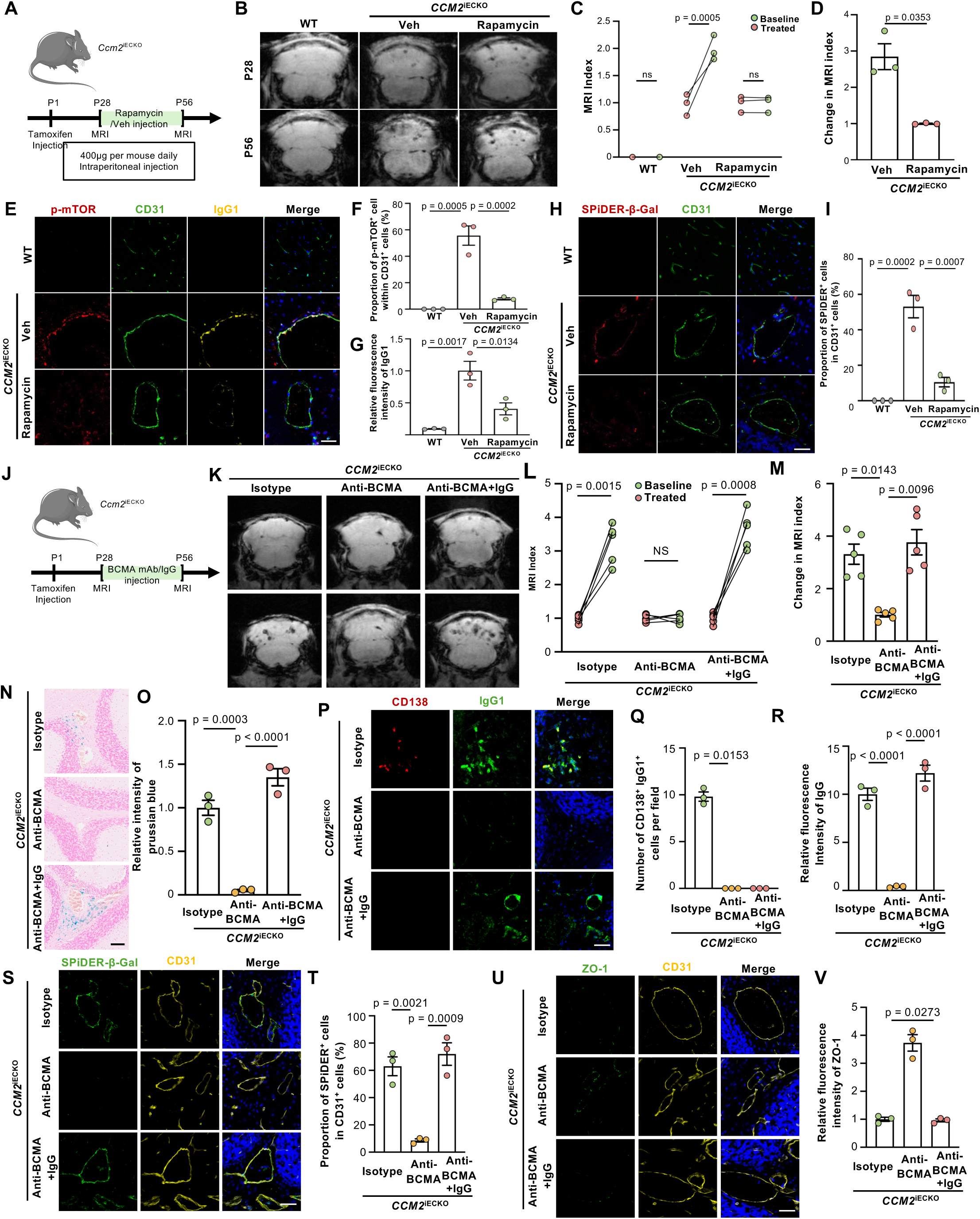
Rapamycin and BCMA-targeted plasma cell intervention attenuate IgG-associated endothelial senescence and lesion progression in *Ccm2*^iECKO^ mice. **(A)** Schematic workflow of Rapamycin treatment in *Ccm2*^iECKO^ mice. Mice underwent MRI examinations at 4 and 8 weeks of age. Rapamycin was administered intraperitoneally at a dose of 400 μg daily. **(B)** Representative MRI scans before and after treatment. **(C)** Quantification of MRI index of mice pre-and post-treatment rapamycin treated or untreated. n = 3. Statistical analysis was performed using paired t-test. **(D)** Relative fold change in MRI index normalized to baseline. Data are presented as mean ± SEM (n = 3 per group). Statistical analysis was performed using unpaired t-test. **(E)** Co-staining of CD31, p-mTOR and IgG1 demonstrating IgG1 deposition and mTOR activation of endothelial cells in rapamycin treated and untreated mice. Scale bar = 50 μm. **(F)** Quantification of the proportion of p-mTOR^+^ cells in endothelial cells. Data are presented as mean ± SEM (n = 3 per group). Three randomly selected fields were analyzed to represent each sample. Statistical analysis was performed using unpaired t-test. **(G)** Quantification of the relative fluorescence intensity of IgG1. Data are presented as mean ± SEM (n = 3 per group). Three randomly selected fields were analyzed to represent each sample. Statistical analysis was performed using unpaired t-test. Relative to Veh group. **(H)** Co-staining of SPiDER-β-Gal and CD31 demonstrating endothelial senescence in rapamycin treated and untreated mice. Scale bar = 50 μm. **(I)** Quantification of the proportion of SPiDER-positive cells in endothelial cells. Data are presented as mean ± SEM (n = 3 per group). Three randomly selected fields were analyzed to represent each sample. Statistical analysis was performed using unpaired t-test. Relative to Veh group. **(J)** Schematic workflow of BCMA antibody treatment in *Ccm2*^iECKO^ mice. Mice underwent MRI examinations at 4 and 8 weeks of age. **(K)** Representative MRI scans before and after treatment. **(L)** Quantification of MRI index of mice pre-and post-treatment. n = 5. Statistical analysis was performed using paired t-test. **(M)** Relative fold change in MRI index normalized to baseline. Data are presented as mean ± SEM (n = 5 per group). Statistical analysis was performed using one-way ANOVA. **(N)** Representative images of Prussian blue staining of perilesional hemorrhage in mice. Scale bar = 100 μm. **(O)** Quantification of Prussian blue staining of BCMA mAb or IgG treated and untreated mice. Data are presented as mean ± SEM (n = 3 per group). Three randomly selected fields were analyzed to represent each sample. Statistical analysis was performed using one-way ANOVA. Relative to isotype control group. **(P)** Representative staining for CD138 plasma cells and IgG cells in mice. Scale bar = 30 μm. **(Q)** Quantification of CD138 IgG plasma cells following treatment. Data are presented as mean ± SEM (n = 3 per group). Three randomly selected fields were analyzed to represent each sample. Statistical analysis was performed using one-way ANOVA. **(R)** Quantification of IgG following treatment. Data are presented as mean ± SEM (n = 3 per group). Three randomly selected fields were analyzed to represent each sample. Statistical analysis was performed using one-way ANOVA. **(S)** Co-staining of SPiDER-β-Gal and CD31 demonstrating the senescence of endothelial cells in treated and control mice. Scale bar = 50 μm. **(T)** Quantification of the proportion of senescent endothelial cells in anti-BCMA ± IgG rescue mice. Data are presented as mean ± SEM (n = 3 per group). Three randomly selected fields were analyzed to represent each sample. Statistical analysis was performed using one-way ANOVA. **(U)** Co-staining of CD31 and the tight junction marker ZO-1 showing endothelial barrier integrity after treatment. Scale bar = 50 μm. **(V)** Quantification of the relative fluorescence intensity of ZO-1. Data are presented as mean ± SEM (n = 3 per group). Three randomly selected fields were analyzed to represent each sample. Statistical analysis was performed using one-way ANOVA. Relative to isotype treated group.

### Plasma cell depletion and IgG rescue establish IgG as a pathogenic effector in CCM

We next tested whether plasma cell-derived IgG directly contributes to CCM progression by depleting plasma cells with an anti-BCMA monoclonal antibody, with or without exogenous native IgG supplementation **(Figure 6J)**. Anti-BCMA monoclonal antibody treatment reduced MRI-based lesion index and stabilized lesion burden over time. In contrast, mice receiving anti-BCMA monoclonal antibody together with exogenous native IgG supplementation showed a restoration of lesion progression **(Figures 6K-M)**.

Histological analysis further confirmed the effects of BCMA-targeted plasma cell depletion and IgG rescue. Anti-BCMA treatment markedly reduced CD138^+^ plasma cells, lesional IgG deposition, perilesional hemorrhage, and endothelial senescence while restoring endothelial ZO-1 expression. In contrast, native IgG supplementation restored lesional IgG accumulation and endothelial senescence and reversed the recovery of ZO-1 expression without replenishing plasma cells **(Figures 6N-V),** demonstrating that IgG is a key downstream effector of plasma cell–mediated endothelial senescence and disease progression. To confirm the biological activity of murine native IgG, mouse endothelial cells were treated with purified murine IgG in vitro. Murine IgG increased NF-κB activation compared with vehicle-treated cells **(Figure S12A)**, consistent with our findings in human endothelial cells. Together, these results demonstrate that plasma cell-derived IgG promotes endothelial NF-κB activation, senescence, and CCM progression in vivo, and that interrupting the plasma cell–IgG axis ameliorates disease.

### Anti-CD38 therapy attenuates IgG-associated vascular dysfunction in CCMs

Having established a causal role for plasma cell–derived IgG using BCMA-mediated plasma cell depletion, we next sought to evaluate a more clinically translatable therapeutic strategy. Although BCMA served as a mechanistic tool to demonstrate the contribution of plasma cells to CCM pathogenesis, neurotoxicity is a recognized concern across anti-BCMA therapies for clinical translation ^86^. In contrast, anti-CD38 monoclonal antibodies (mAb) are established plasma cell– depleting therapies with expanding use in autoimmune diseases, including neuroimmunologic disorders ^87–89^, making CD38 an attractive therapeutic target for CCM.

In CCM lesions, CD38 was abundantly expressed by plasma cells and was present at significantly higher levels than in circulating plasma cells from peripheral blood **(Figures 7A and S12B)**. IHC analysis further showed substantial overlap between CD38 and the plasma cell marker CD138 within CCM lesions **(Figure 7B)**.

**Figure 7.**
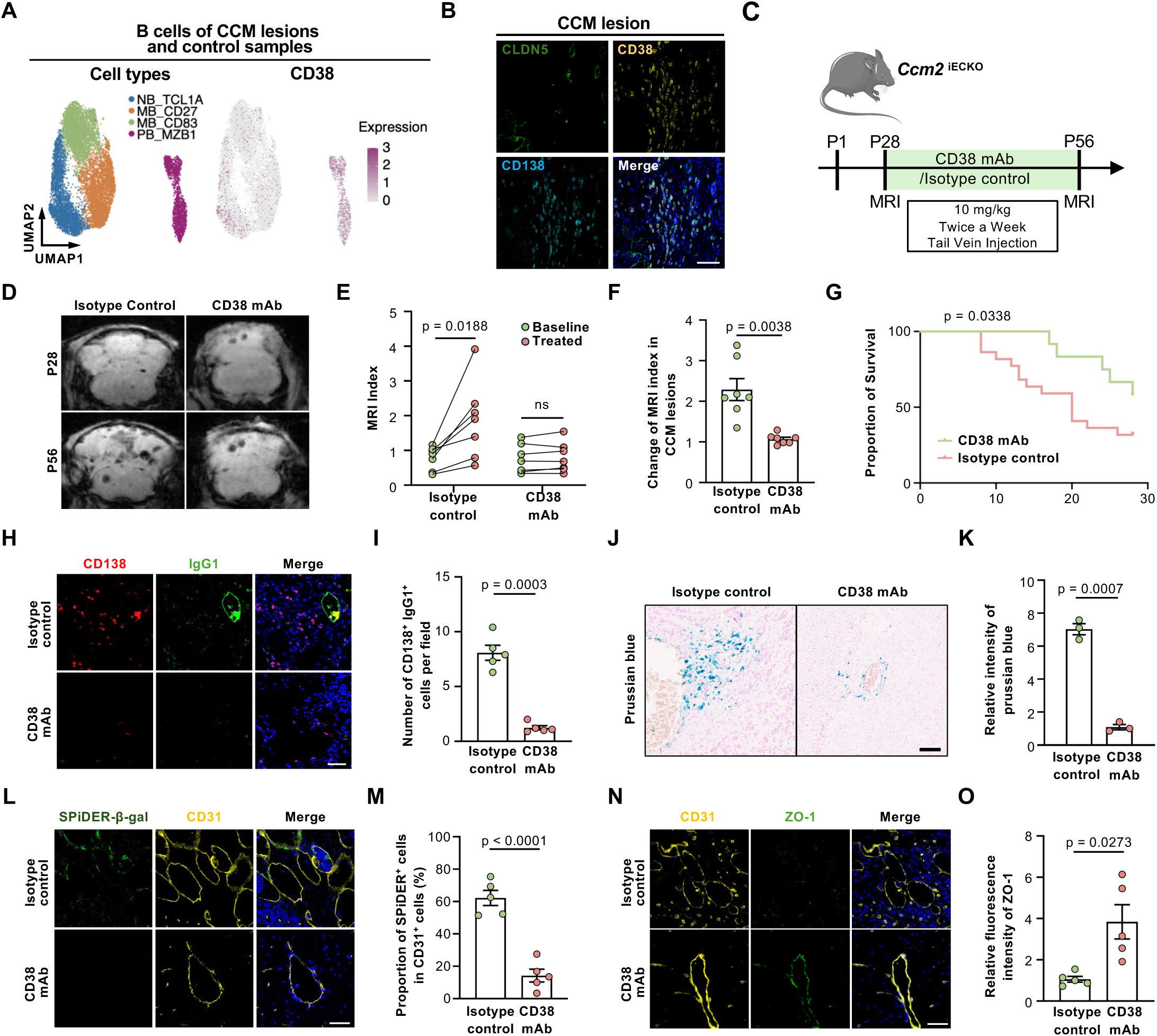
CD38 as a therapeutic target for CCM reverses plasma cell IgG–induced endothelial senescence. **(A)** UMAP visualization showing CD38 expression in B cells. **(B)** Representative immunofluorescence images showing spatial overlap of CD38 with the plasma cell marker CD138. Scale bar = 50 μm. **(C)** Schematic workflow of CD38 antibody treatment in *Ccm2*^iECKO^ mice. Mice underwent MRI examinations at 4 and 8 weeks of age. CD38 monoclonal antibody was administered intravenously via the tail vein at a dose of 10 mg/kg, twice per week. **(D)** Representative MRI images before and after treatment. **(E)** Quantification of MRI index of mice pre-and post-treatment (n = 7). Statistical analysis was performed using paired t-test. **(F)** Relative fold change in MRI index normalized to baseline. Data are presented as mean ± SEM (n = 7 per group). Statistical analysis was performed using unpaired t-test. **(G)** Kaplan–Meier survival curves comparing CD38 mAb treated and control mice. **(H)** Representative staining for CD138 plasma cells and IgG1 cells in CD38 mAb treated and untreated mice. Scale bar = 30 μm. **(I)** Quantification of IgG1^+^ plasma cell following treatment. Data are presented as mean ± SEM (n = 5 per group). Three randomly selected fields were analyzed to represent each sample. Statistical analysis was performed using unpaired t-test. **(J)** Representative images of Prussian blue staining of perilesional hemorrhage in CD38 mAb treated and untreated mice. Scale bar = 100 μm. **(K)** Quantification of Prussian blue staining of CD38 mAb treated and untreated mice. Data are presented as mean ± SEM (n = 3 per group). Three randomly selected fields were analyzed to represent each sample. Statistical analysis was performed using unpaired t-test. Relative to isotype control group. **(L)** Co-staining of SPiDER-β-Gal and CD31 demonstrating the senescence of endothelial cells in treated and control mice. Scale bar = 50 μm. **(M)** Quantification of the proportion of senescent endothelial cells in CD38 mAb treated and untreated mice. Data are presented as mean ± SEM (n = 5 per group). Three randomly selected fields were analyzed to represent each sample. Statistical analysis was performed using unpaired t-test. **(N)** Co-staining of CD31 and the tight junction marker ZO-1 showing endothelial barrier integrity after treatment. Scale bar = 50 μm. **(O)** Quantification of the relative fluorescence intensity of ZO-1. Data are presented as mean ± SEM (n = 5 per group). Three randomly selected fields were analyzed to represent each sample. Statistical analysis was performed using unpaired t-test. Relative to isotype treated group.

To determine whether CD38-targeted plasma cell depletion could mitigate CCM pathogenesis, CD38 mAb was administered by tail-vein injection to 4-week-old CCM mice, and lesion progression was evaluated 4 weeks later **(Figure 7C)**. Compared to control, CD38 mAb-treated mice prevented lesion progression over time **(Figures 7D-F)** and significantly prolonged survival **(Figure 7G)**.

CD38 mAb treatment markedly reduc**ed** CD138 IgG1 plasma cells **(Figures 7H–I)**. Consistently, the number of CD38 cells within CCM lesions was also reduced **(Figures S12C–D)**. Prussian blue staining revealed reduced perilesional hemorrhage after CD38 mAb treatment **(Figures 7J-K)**. SPiDER-β-Gal staining showed reduced senescent endothelial cells **(Figures 7L-M)**, whereas ZO-1 immunostaining showed restoration of endothelial junctional integrity **(Figures 7N-O)**. Together, these findings demonstrate that targeting plasma cells with a clinically established anti-CD38 therapy reduces pathogenic IgG production, mitigates endothelial dysfunction, and slows CCM progression, supporting plasma cell depletion as a promising therapeutic strategy for CCM.

## DISCUSSION

In this study, we integrated single-cell and spatial transcriptomics with mechanistic and therapeutic studies to define a plasma cell–endothelial axis that contributes to CCM progression. Our findings extend previous histological observations of B-cell infiltration ^50,53–55^ by identifying lysosomal dysfunction-mediated IgG accumulation as a pathogenic contributor to endothelial senescence and by implicating an mTOR–lysosome–NF-κB axis in the interaction between humoral immunity and endothelial dysfunction.

These findings extend emerging evidence that IgG contributes to age-associated tissue dysfunction beyond myeloid cells. Previous studies implicated IgG in senescence-associated phenotypes in macrophages and microglia ^11^, whereas our data demonstrate that plasma cell-derived IgG can directly induce endothelial senescence through NF-κB signaling. In parallel, mTOR-dependent lysosomal dysfunction promotes intracellular IgG accumulation and enhances endothelial susceptibility to IgG-associated injury. Thus, CCM genetic alterations may establish a state of endothelial vulnerability in which humoral immune signals amplify vascular dysfunction.

mTOR is a central regulator of cellular metabolism and homeostasis, coordinating lysosomal function, immune signaling, protein synthesis, autophagy, redox balance, and mitochondrial activity ^90^. Dysregulated mTOR signaling has been implicated in vascular disease, including CCM ^79,83,91^. In our study, rapamycin reduced endothelial IgG accumulation and markedly attenuated endothelial senescence both in vitro and in vivo, supporting lysosomal dysfunction as an important component of mTOR-associated endothelial injury. However, given the pleiotropic functions of mTOR and the broad biological effects of rapamycin, lysosomal dysfunction is unlikely to represent the sole downstream mechanism. Additional processes, including inflammatory signaling, proteostasis, oxidative stress, and mitochondrial metabolism, may also contribute to endothelial dysfunction in CCM and warrant further investigation.

Circulating IgG under physiological conditions is unlikely to be sufficient to induce widespread vascular senescence. Endothelial cells normally maintain intracellular IgG homeostasis through FcRn-dependent, pH-sensitive recycling and lysosomal processing, thereby limiting excessive intracellular IgG accumulation ^92^. In CCM, constitutive mTOR activation impairs lysosomal acidification and IgG handling, favoring intracellular IgG persistence. Together with plasma cell infiltration and blood– brain barrier disruption, these abnormalities increase local IgG availability and create a permissive microenvironment in which IgG acts as a secondary pathogenic signal that amplifies endothelial senescence and lesion progression.

Our study also has important therapeutic implications. Previous work demonstrated that pan-B-cell depletion stabilizes CCM lesions in preclinical models ^53^. Our data further identify plasma cells as a functionally important effector population and show that BCMA-mediated plasma cell depletion, together with IgG rescue, supports a causal role for plasma cell-derived IgG in disease progression. Building on this mechanistic insight, we further demonstrate that anti-CD38 treatment reduces IgG deposition, endothelial senescence, hemorrhage, lesion progression, and mortality in CCM mice, providing a rationale for further evaluation of plasma cell-targeted therapy in CCM.

Several questions remain for future investigation. The antigen specificity, clonality, and subclass distribution of CCM-associated IgG remain incompletely defined. In addition, the molecular mechanisms governing endothelial IgG uptake, intracellular trafficking, Fc-receptor engagement, and lysosomal processing require further investigation. Finally, because CD38 is expressed by multiple immune-cell populations ^93^, contributions from non-plasma-cell targets to the therapeutic effects of CD38-targeted treatment cannot be excluded.

In summary, our findings identify lysosomal dysfunction-mediated IgG as a pathogenic trigger of endothelial senescence and dysfunction in CCM and support selective plasma cell targeting as a potential therapeutic strategy. More broadly, they reveal how genetic endothelial defects and adaptive immunity converge to promote vascular senescence, with potential implications for other inflammatory cerebrovascular diseases.

## RESOURCE AVAILABILITY

### Materials availability

This study did not generate new unique reagents.

### Data and code availability

All raw human sequencing data generated in this study have been deposited in the Genome Sequence Archive for Human (GSA-Human) at the National Genomics Data Center, China National Center for Bioinformation/Beijing Institute of Genomics, Chinese Academy of Sciences, under accession numbers **HRA018293, HRA018294, HRA018295, HRA018296, HRA018297, and HRA019381**. Raw single-cell RNA-sequencing data from the CCM mouse models have been deposited in the Genome Sequence Archive under accession number **CRA042693**. Microscopic images corresponding to the spatial transcriptomic dataset are available in OMIX under accession number **OMIX018213**. All datasets are publicly accessible.

Previously published single cell RNA-seq and bulk RNA-seq analyzed in this study can be found in Supplementary Table 2. Source data are provided with this paper. Custom code used in this study is publicly available at: https://github.com/sheenaseven/CCM-PB. Patient information is available in the supplemental information. Any additional information required to reanalyze the data reported in this paper is available from the lead contact upon request.

## ACKNOWLEDGMENTS

Research in the Cao lab was funded by the NSFC International Cooperation and Exchanges Program (W2411067) and the NSFC General Program (82171267). Research in the Wang lab was supported by the RGC grant (16102522), the ITC grant (ITCPD/17-9), and was also supported by Padma Harilela Professorship and Sinovac Fellowship.

## AUTHOR CONTRIBUTIONS

Y.Cao, R.O.L., J.W., and W.S.H. conceptualized and supervised the study. Y.S., S.Z., R.H., and Z.X. designed and conducted the experiments. R.H. collected the clinical data and human tissue samples. Y.Y. performed the majority of the multi-omics data analyses. Y.S., S.Z., Z.X., and Q.Zhang analyzed the experimental data. H.W. contributed to some of the spatial data analyses. Q.Zhou, Y.Chen, R.S., L.D., and Z.G. contributed to data interpretation. Y.Y., Y.S., and S.Z. wrote the manuscript, which was then revised and proofread by all authors.

## SUPPLEMENTAL INFORMATION

Document S1: Key resources table and Methods, supplemental references

Document S2: Figures S1-S12

Supplementary table: Table S1-S5

