## Supplementary material for "Lysosomal Dysfunction–Mediated IgG Accumulation Promotes Endothelial Senescence and Lesion Progression in Cerebral Cavernous Malformations": CD38_Figures.pdf

### Figure 1

A

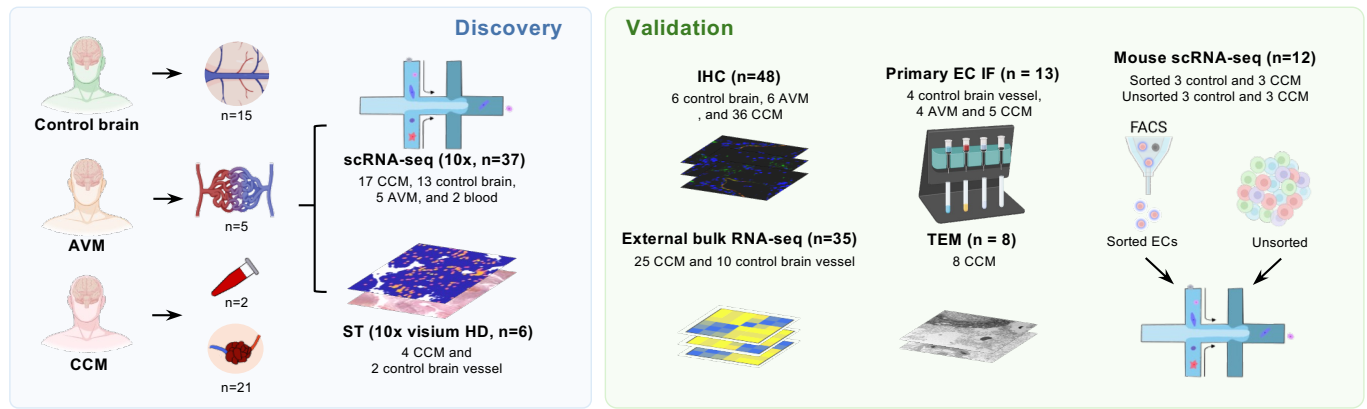

B

#### Major cell types of CCM lesions and control samples

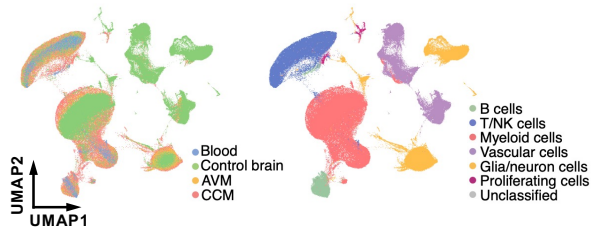

C

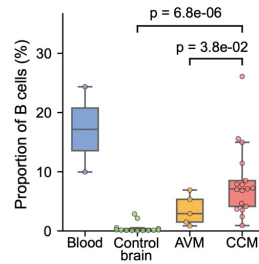

D

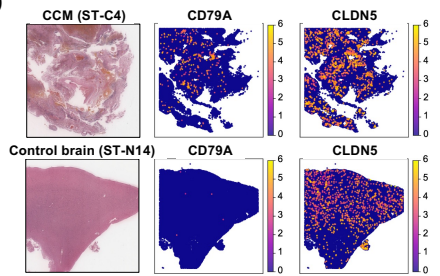

E

#### B cell types of CCM lesions and control samples

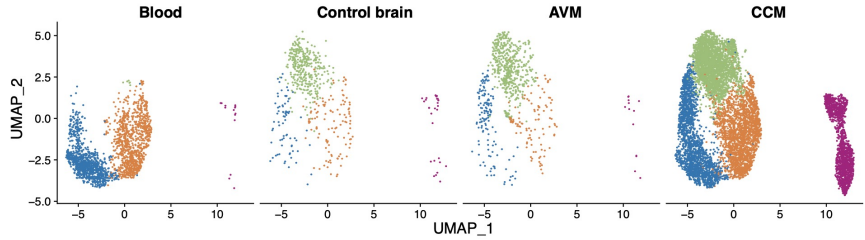

G

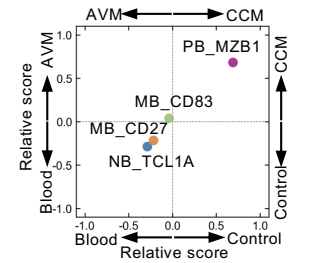

F

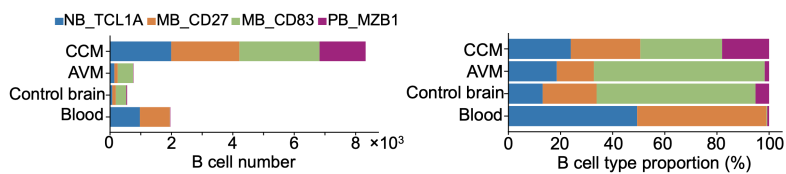

H

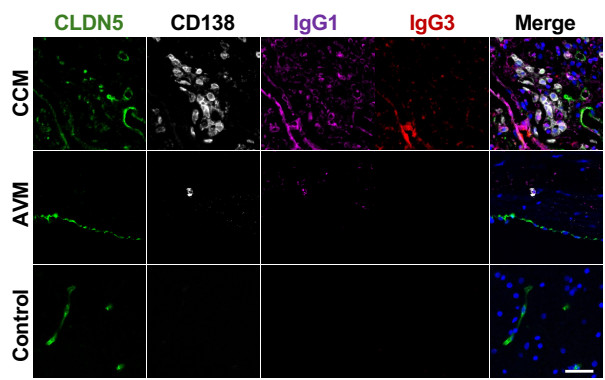

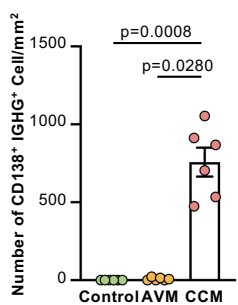

I

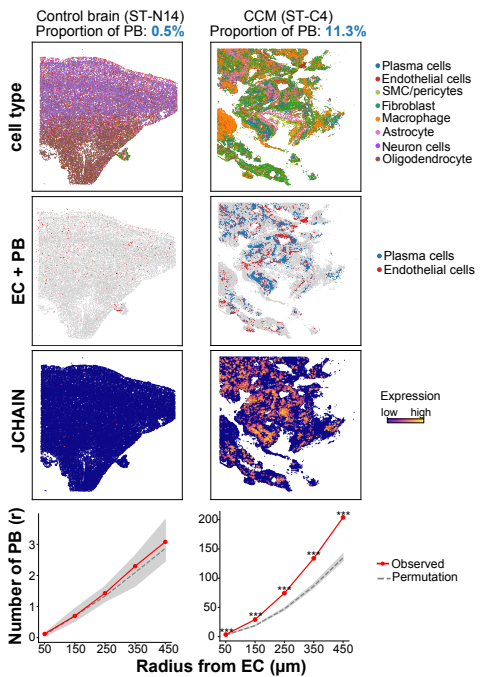

**A**

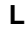

**Figure 3**

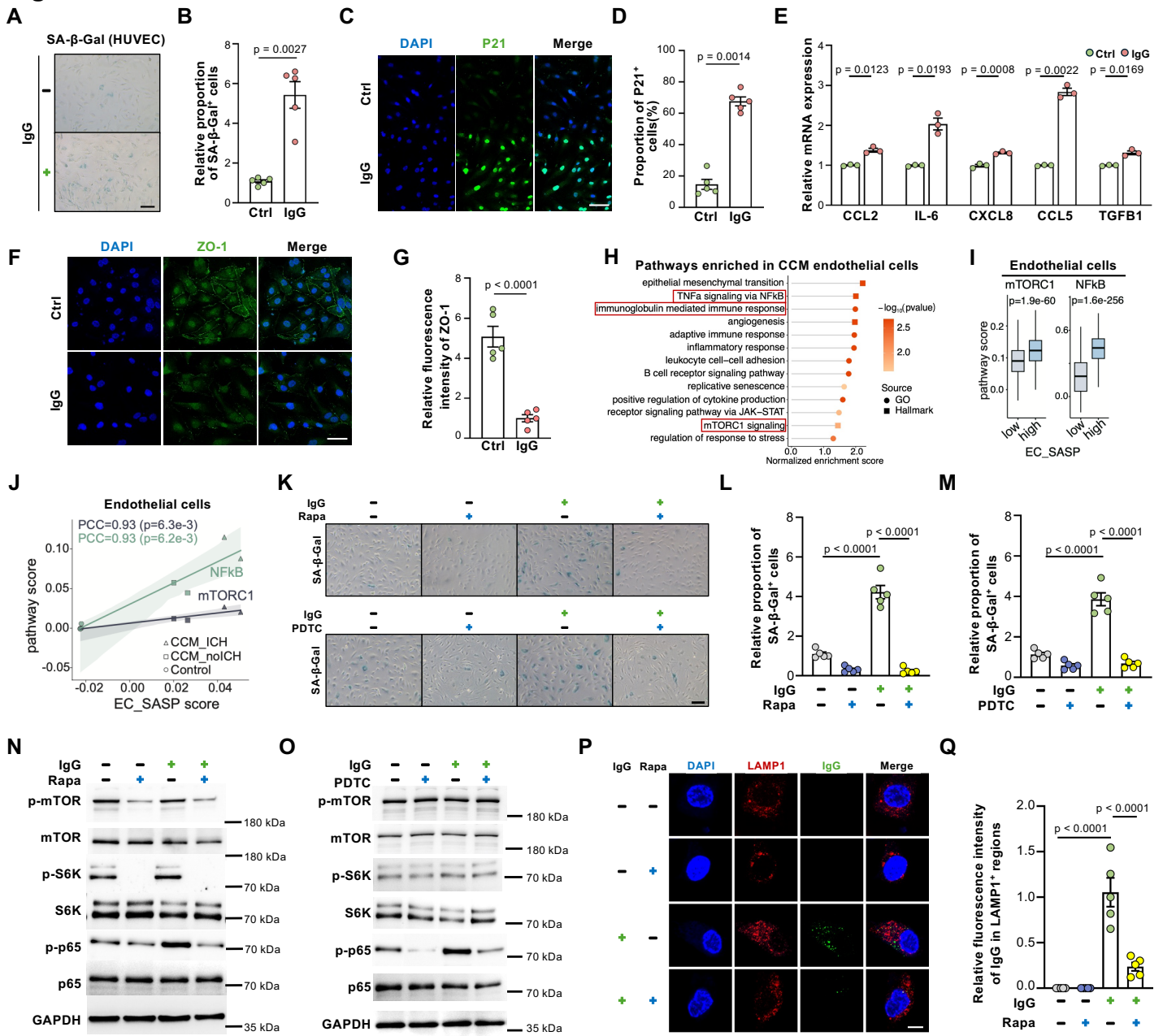

Figure 4

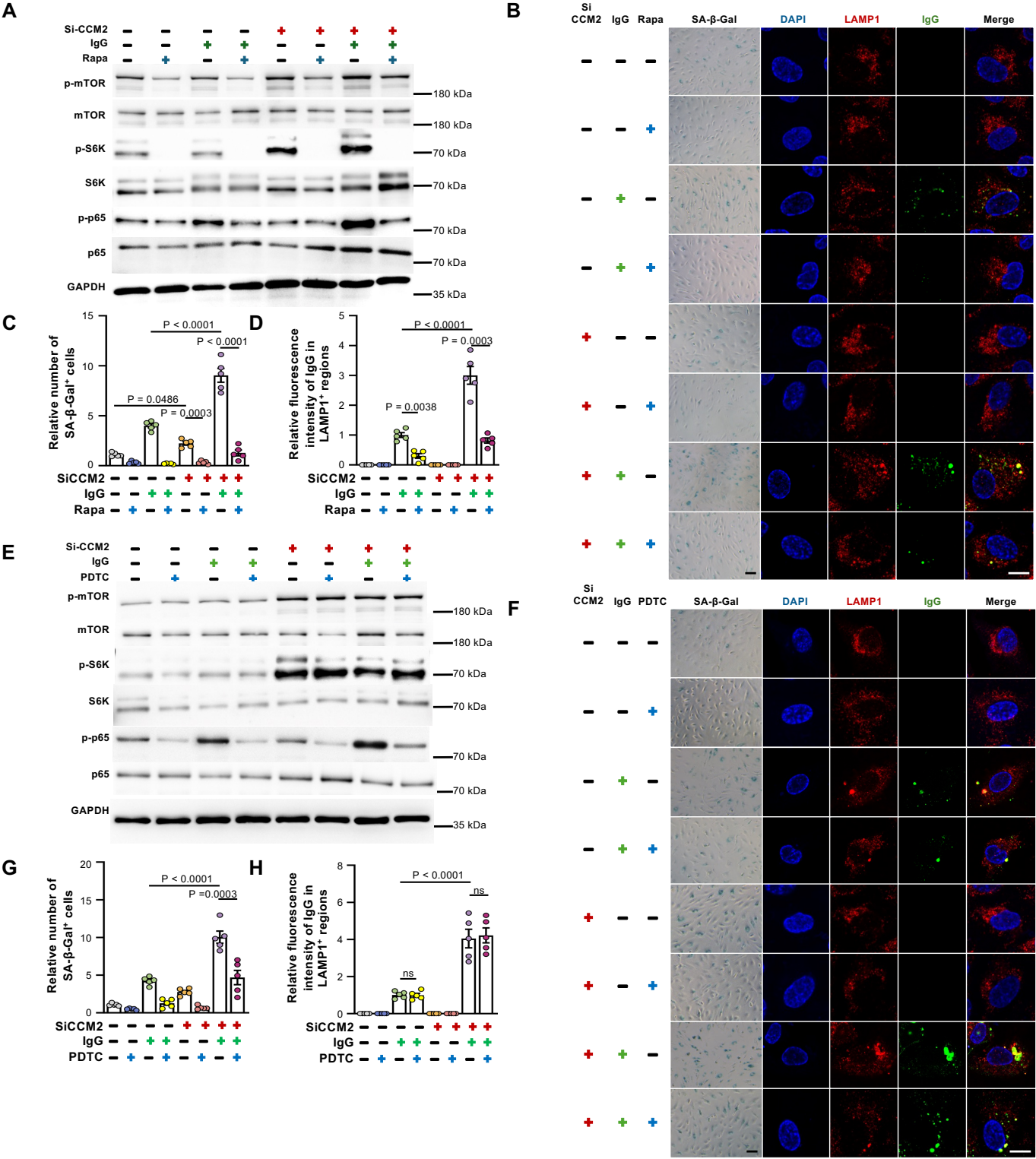

**Figure 5**

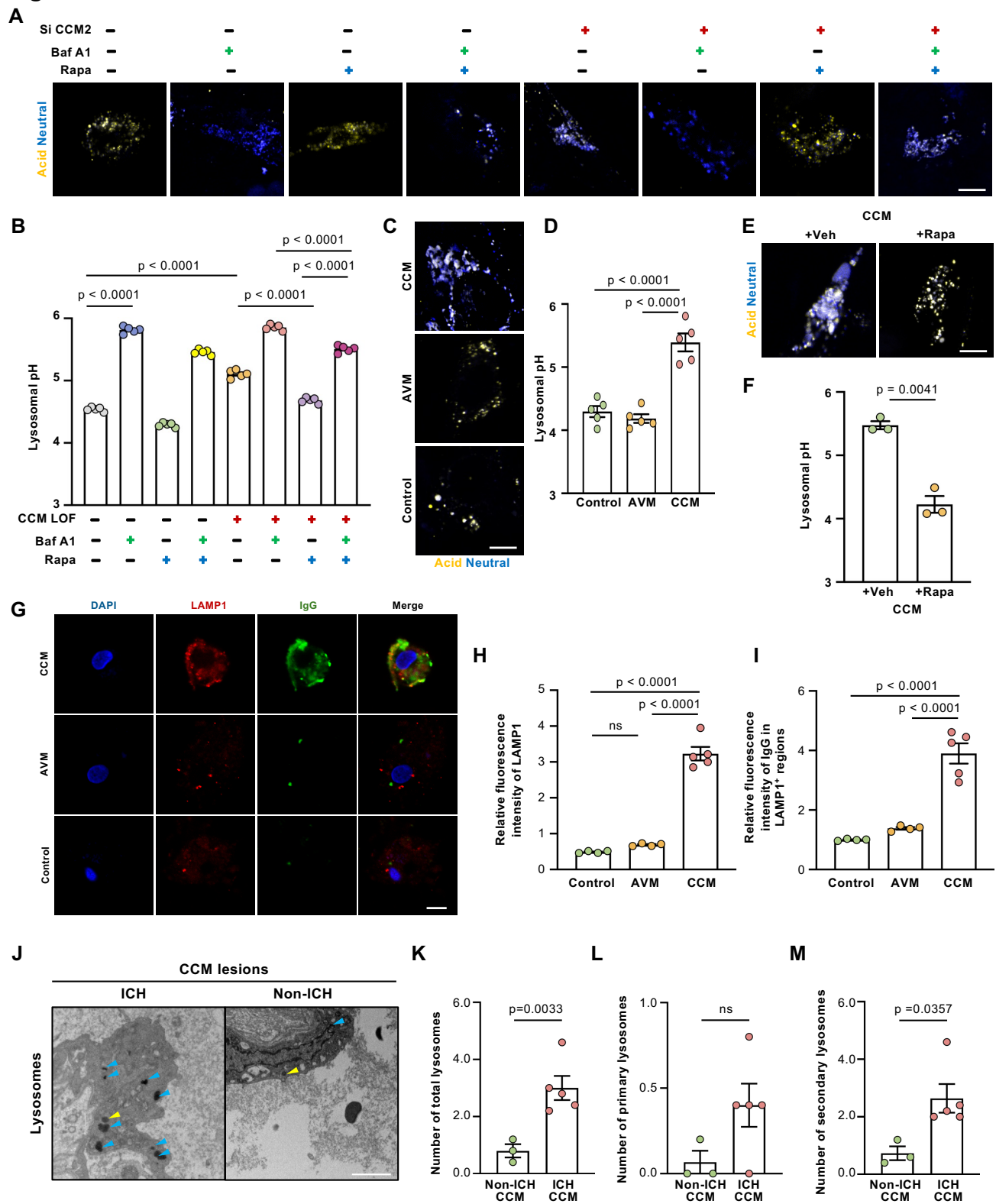

**Figure 6**

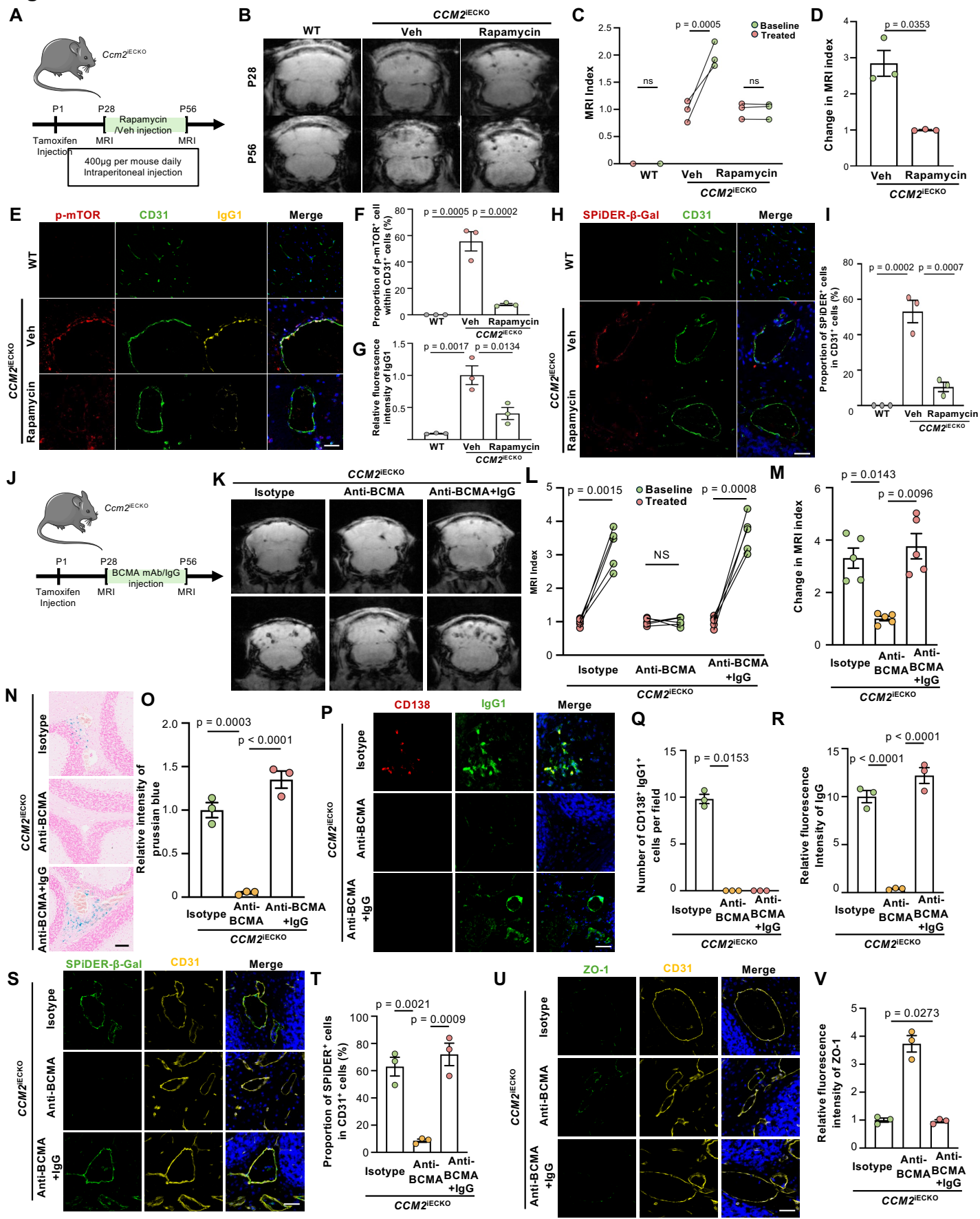

**Figure 7**

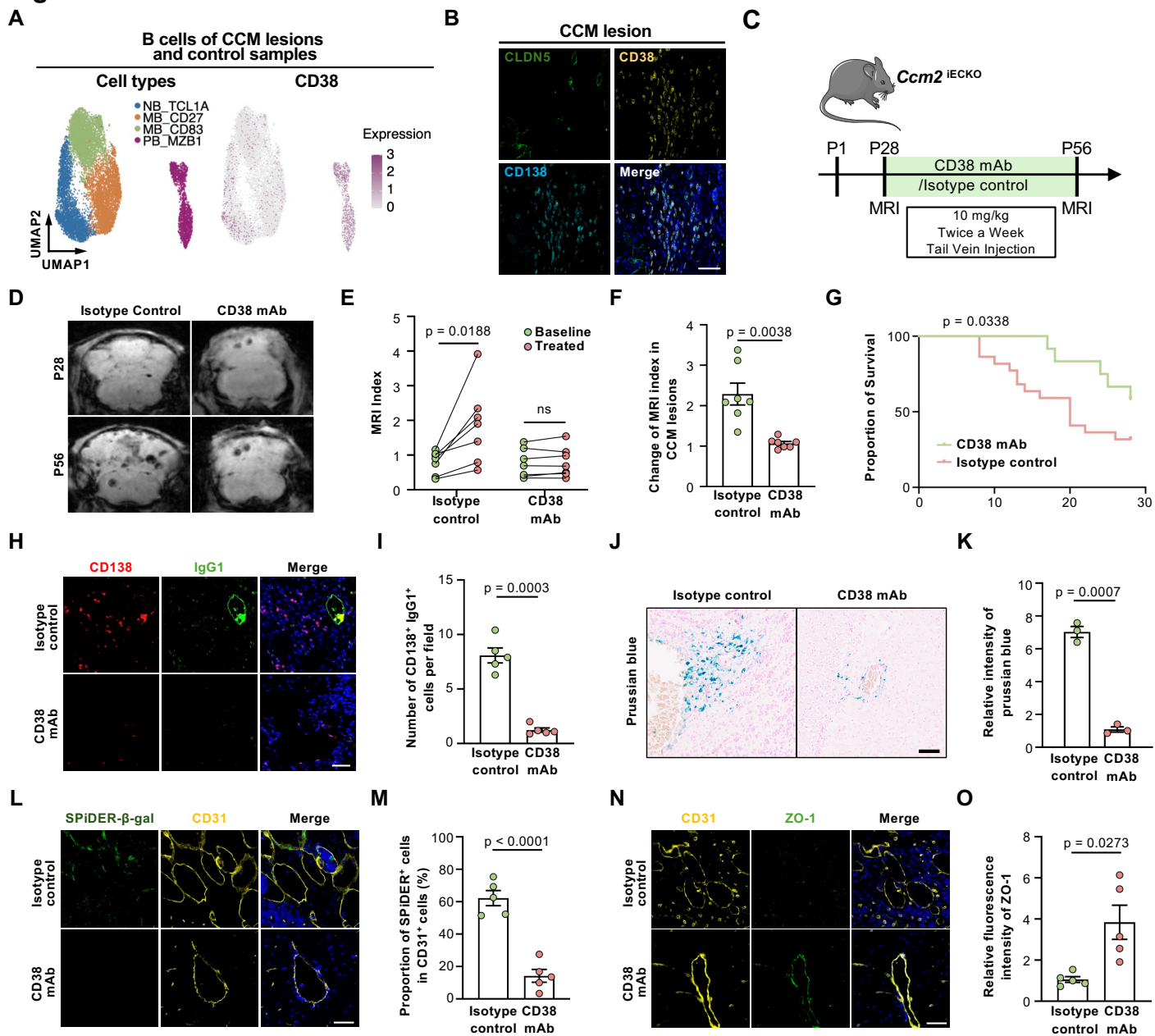
