## Supplementary material for "Lysosomal Dysfunction–Mediated IgG Accumulation Promotes Endothelial Senescence and Lesion Progression in Cerebral Cavernous Malformations": Supplemental_Methods.docx

**Primary and Secondary Antibodies**

| **Antibodies** | **SOURCE** | **IDENTIFIER** |
| --- | --- | --- |
| Rabbit anti-Human CLDN5 | Abcam | Cat#ab15106  RRID: AB_301652 |
| Rabbit anti-Human IgG3 | Abcam | Cat#ab193172 |
| Rabbit anti-Human/Mouse IgG1 | Immunoway | Cat#YT2293 |
| Mouse anti-Human Syndecan1 (CD138) | Abcam | Cat#ab128936  RRID: AB_11150990 |
| Rabbit anti-Mouse CD138 | Thermo | Cat#36-2900 |
| Rabbit anti-Human CD38 | Abcam | Cat#ab108403  RRID: AB_10890803 |
| Rabbit anti-Mouse CD38 | Cell signaling technology | Cat#68336 |
| Rabbit anti-Human GAPDH | Cell signaling technology | Cat#2118  RRID: AB_561053 |
| Rabbit anti-Human/Mouse ZO-1 (IHC) | Abcam | Cat#ab221547  RRID: AB_2892660 |
| Rabbit anti-Human P21 (IF) | Cell signaling technology | Cat#2947  RRID: AB_823586 |
| Rabbit anti-Human ZO-1 (IF) | Cell signaling technology | Cat#13663  RRID: AB_2798287 |
| Mouse anti-Human LAMP1 (IF) | Abcam | Cat#ab25630  RRID: AB_470708 |
| Rabbit anti-Human Phospho-mTOR (Ser2448) (WB) | Cell signaling technology | Cat#5536  RRID: AB_10691552 |
| Rabbit anti-Human mTOR (WB) | Cell signaling technology | Cat#2983  RRID: AB_2105622 |
| Rabbit anti-Human Phospho-P65 (Ser536) (WB) | Cell signaling technology | Cat#3033  RRID: AB_331284 |
| Rabbit anti-Human P65 (WB) | Cell signaling technology | Cat#8242  RRID: AB_10859369 |
| Rabbit anti-Human Phospho-p70 S6 Kinase (Thr389) (WB) | Cell signaling technology | Cat#9234  RRID: AB_2269803 |
| Rabbit anti-Human P70 S6 Kinase (WB) | Cell signaling technology | Cat#2708  RRID: AB_390722 |
| Rabbit anti-Mouse P21 (IHC) | Abcam | Cat#ab188224  RRID: AB_2734729 |
| Rabbit anti-Mouse CD31 (IHC) | Cell signaling technology | Cat#77699  RRID: AB_2722705 |
| PE rat anti-mouse CD31 | Thermo Fisher | Cat#12-0311-82  RRID: AB_465632 |
| FITC rat anti-mouse CD45 | Thermo Fisher | Cat#11-0451-82  RRID: AB_465050 |
| Human native IgG | Abcam | Ab91102 |
| Mouse native IgG | Abcam | Ab198772 |
| Anti-mouse CD38 | Bio X cell | CP052 |
| mouse IgG2a isotype control | Bio X cell | BE0085 |
| anti-mouse CD269 (BCMA) Antibody | BioLegned | 943604  RRID: AB_2892507 |
| IgG2a, κ Isotype Ctrl Antibody | BioLegned | 400565  RRID: AB_11147167 |
| Goat anti-Rabbit IgG-HRP | Abmart | Cat#M21002 |
| Goat anti-Mouse IgG HRP | Abmart | Cat#M21001 |
| Goat anti-Human IgG H&L (DyLight® 488) | Abcam | Cat#ab96907  RRID: AB_10680176 |
| Goat anti-Mouse IgG H&L (Alexa Fluor® 594) | Abcam | Cat#ab150116  RRID: AB_2650601 |

**Datasets Analyzed in This Study**

| **Deposited Data** | **SOURCE** | **IDENTIFIER** |
| --- | --- | --- |
| Single-cell RNA-sequencing data | This study |  |
| Visium HD ST data | This study |  |
| External scRNA-seq data of controls | Wälchli et al. ^94^ |  |
| External scRNA-seq data of controls and CCMs | Han et al. ^95^ | GEO: GSE294555 |
| Bulk RNA-seq data of controls and CCMs | Subhash et al. ^96^; Lyne et al. ^97^; Koskimäki et al. ^98^ | GEO: GSE137596; GSE130174; GSE123968 |
| Whole-brain mouse scRNA-seq data | This study |  |
| EC–sorted mouse scRNA-seq data | This study |  |
| external EC–sorted mouse scRNA-seq data | Orsenigo et al. ^99^ | GEO: GSE155788 |

**Software and Packages Used for Bioinformatics Analyses**

| **Software and Algorithms** | | **SOURCE** | **IDENTIFIER** |
| --- | --- | --- | --- |
| Cellranger (v5.0.1; v7.0.1) | 10x Genomics | | https://www.10xgenomics.com/support/software/cell-ranger/latest |
| Seurat (v4.3.0) | Hao et al. ^100^ | | https://satijalab.org/seurat |
| scDblFinder (v1.12.0) | Germain et al. ^101^ | | https://github.com/plger/scDblFinder |
| Harmony (v1.2.3) | Korsunsky et al. ^102^ | | https://github.com/immunogenomics/harmony |
| Garnett (v0.2.19) | Pliner et al. ^103^ | | https://cole-trapnell-lab.github.io/garnett |
| clusterProfiler (4.10.0) | Yu et al. ^104^ | | https://github.com/YuLab-SMU/clusterProfiler |
| Monocle3 (v1.3.1) | Qiu et al. ^105^ | | https://github.com/cole-trapnell-lab/monocle3 |
| scanpy (v1.9.1) | Wolf et al. ^106^ | | https://github.com/scverse/scanpy |
| scipy (v1.7.3) | Virtanen et al. ^107^ | | https://github.com/scipy/scipy |

**METHOD DETAILS**

**Experimental Model and Subject Details**

**Tissue samples/Patients**

We collected CCMs or AVMs lesions from patients who were clinically diagnosed by medical history, neuroradiological examination (CT, MRI and/or DSA) and the standard pathological results. Non-lesional specimens were obtained from patients undergoing focal resection for epilepsy. The collection of human samples was approved by all patients under written informed consent. This study was conducted in accordance with the Declaration of Helsinki and was approved by the Internal Review and Ethics Committee of the Beijing Tiantan Hospital. For radiological classification, two board-certified neurosurgeons independently evaluated the MRI images. Concordant assessments were accepted as the final classification. In the event of a disagreement, a third senior neurosurgeon adjudicated the final grade.

**Mouse studies**

The Animal Welfare and Ethics Committee of Beijing Neurosurgical Institute Laboratory approved all animal ethics and protocols. All mice were housed in individual, ventilated cages with 12-hour light/dark cycles, with food and water ad libitum. All animals were housed in a specific pathogen-free environment in a vivarium. The *Ccm2*^fl/fl^ mice were generated on the C57BL/6J background and created by CRISPR/Cas-mediated genome engineering at Cyagen Biosciences (China). After crossing the above mice with *Cdh5*-CreERT2 mice (purchased from Cyagen Biosciences), *Cdh5*-CreERT2; *Ccm2*^fl/fl^ (*Ccm2*^iECKO^) mice were obtained.

For all mouse model experiments, at one-day post-birth (P1), pups were intragastrically injected by 30-gauge needle with 25 μg of 4-hydroxytamoxifen (4-OHT, Sigma Aldrich, H7904) freshly dissolved in a 9% ethanol/corn oil (volume/volume) vehicle (50 μL total volume per injection). Both male and female mice were used. The mouse models were confirmed at 4 weeks by 7.0 T small animal MRI scanner (Bruker, Germany), after which were randomized into different groups. No animals or data points were excluded from the analysis. Investigators were blinded to group allocation during data acquisition and analysis. The experimental unit was a single animal.

**Human single-cell RNA-seq data processing and analysis**

***scRNA-seq data collection***

Published human scRNA-seq vasculature datasets were collected from Wälchli et al. and Han et al. (GSE294555), including external control brains and one CCM lesion used for integrated analysis. The newly generated scRNA-seq dataset comprised 27 samples collected at Beijing Tiantan Hospital. Detailed clinical information is provided in Supplementary Table 1.

***Tissue dissociation and cell purification***

For the newly generated patients’ samples in this study, freshly resected samples were placed at 4 ℃ and transported to the laboratory. Under a dissecting microscope at 2-5× magnification, microsurgical instruments were used to dissect all visible vascular tissues for subsequent sequencing. No differences in vascular isolation methods were applied between normal cerebral cortex, AVM or CCM tissues. Tissues were transported in sterile culture dish with 10 ml 1× Dulbecco's Phosphate-Buffered Saline (DPBS; Thermo Fisher, Cat. no. 14190144) on ice to remove the residual tissue storage solution, then minced on ice. We used dissociation enzyme 0.25% Trypsin (Thermo Fisher, Cat. no. 25200072) and 10 μg/mL DNase I (Sigma, Cat. no. 11284932001) dissolved in PBS (phosphate-buffered saline) (1×PBS, P1020, Solarbio, Beijing) with 5% Fetal Bovine Serum (FBS; Sciencell, Cat. no. 0025) to digest the tissues. Tissues were dissociated at 37 °C with a shaking speed of 50 rpm for about 40 min. We repeatedly collected the dissociated cells at interval of 20 min to increase cell yield and viability. Cell suspensions were filtered using a 40 μm nylon cell strainer and red blood cells were removed by 1× Red Blood Cell Lysis Solution (Thermo Fisher, Cat. no. 00-4333-57). Dissociated cells were washed with 1× DPBS containing 2% FBS. Cells were stained with 0.4% Trypan blue (Thermo Fisher, Cat. no. 15250061) to check the viability on Countess® II Automated Cell Counter (Thermo Fisher). scRNA-seq was performed by Shanghai Biotechnology Corporation (Shanghai, China).

***10x library preparation and sequencing***

Beads with unique molecular identifier (UMI) and cell barcodes were loaded close to saturation, so that each cell was paired with a bead in a Gel Beads-in-emulsion (GEM). After exposure to cell lysis buffer, polyadenylated RNA molecules hybridized to the beads. Beads were retrieved into a single tube for reverse transcription. On cDNA synthesis, each cDNA molecule was tagged on the 5’end (that is, the 3’end of a messenger RNA transcript) with UMI and cell label indicating its cell of origin. Briefly, 10× beads that were then subject to second-strand cDNA synthesis, adaptor ligation, and universal amplification. Sequencing libraries were prepared using randomly interrupted whole-transcriptome amplification products to enrich the 3’ end of the transcripts linked with the cell barcode and UMI. All the remaining procedures including the library construction were performed according to the standard manufacturer’s protocol (CG000206 Rev D). Sequencing libraries were quantified using a High Sensitivity DNA Chip (Agilent) on a Bioanalyzer 2100 and the Qubit High Sensitivity DNA Assay (Thermo Fisher Scientific). The libraries were sequenced on NovaSeq6000 (Illumina) using 2x150 chemistry.

***scRNA-seq data analysis***

Gene expression matrices for the newly enrolled samples were generated using the CellRanger (v5.0.1; 10x Genomics) with the human reference genome (GRCh38-1.2.0). Publicly available and in-house datasets were subsequently merged for joint analysis. We performed quality control filtering via the R package Seurat (v4.3.0)^100^ Cells were removed: (1) UMI counts ≤ 2000 or ≥ 15,000; (2) expressed ≤ 300 genes or ≥ 4,000 genes; (3) ≥ 10% of mitochondrial UMIs; (4) ≥ 1% of hemoglobin UMIs. Genes detected by fewer than 10 cells were excluded. Doublets were identified and excluded using scDblFinder (v1.12.0) ^101^ . The filtered expression matrix was normalized for library size and log-transformed using the “NormalizeData” function, and then genes were scaled to unit variance with the “ScaleData” function. We performed principal component analysis on the top 2,000 variable genes identified using “FindVariableFeatures” function (selection.method=’vst’). Batch effects arising from different sources and donors were corrected using Harmony (v1.2.3)^102^. The Louvain algorithm was used to cluster cells via the functions “FindNeighbors” (k-nearest neighbors = 20) and “FindClusters” (resolution = 0.6). Clusters were then visualized by Uniform Manifold Approximation and Projection.

***Cell type annotation***

The major cell identity was manually annotated by conventional markers described in Fig. S1C. To assess the robustness of clusters, Garnett^103^ (v0.2.19, an automated cell type classification tool) was implemented for distinguishing the 5 major cell types. The gene signatures were defined (*CLDN5*, *VWF*, *COL3A1*, *KCNJ8*, *TAGLN* for vascular cells; *PLP1*, *MBP* for glia cells; *CD14*, *FCGR3A*, *LYZ*, *CD68*, *CD163*, *C1QA*, *CD1C*, *FCER1A*, *IL3RA*, *TPSB2* for myeloid lineage cells; *CD19*, *MS4A1*, *CD79A* for B cells; *CD3D*, *CD3E*, *CD3G*, *GNLY*, *NKG7* for T cells). We employed the *train_cell_classifier* function with the org.Hs.eg.db gene database and default parameters to train the models. The *classify_cells* function (cluster_extend = TRUE) was used to check the consistency between Garnett-predicted cell type labels with the manual annotations.

***Tissue enrichment analysis***

The tissue enrichment analysis was developed to systematically assess and visualize the relative distribution of cell clusters across different tissue groups. This approach utilized a two-dimensional representation, assigning all cell clusters to four distinct quadrants based on the median cluster proportions (MCP) calculated for each group.

Quadrant I: CCM-enriched clusters with x axis defined as *MCP_CCM_ - max(MCP_blood_, MCP_AVM_)* and y axis defined as *MCP_CCM_ - max(MCP_blood_, MCP_control_*).

Quadrant II: AVM-enriched clusters with x axis defined as *max(MCP_CCM_, MCP_control_) - MCP_AVM_* and y axis defined as *MCP_AVM_ - max(MCP_blood_, MCP_control_)*.

Quadrant III: blood-enriched clusters with x axis defined as *max(MCP_CCM_, MCP_control_) - MCP_blood_* and y axis defined as *max(MCP_AVM_, MCP_CCM_) - MCP_blood_*.

Quadrant IV: control-enriched clusters with x axis defined as MCPcontrol - max(MCPAVM, MCPblood) and y axis defined as max(MCPCCM, MCPAVM) - MCPcontrol.

***Differential-expression analysis and pathway enrichment analysis***

Differentially expressed genes (DEGs) between CCM and control samples were identified using Seurat FindMarkers (logfc.threshold = 0, min.pct = 0, Wilcoxon rank-sum test with Bonferroni correction). Marker genes distinguishing B-cell subsets were identified using FindAllMarkers. GO enrichment and GSEA were performed using clusterProfiler, with hallmark gene sets retrieved from MSigDB. mTORC1 and NF-κB pathway scores were computed using Seurat AddModuleScore.

***Endothelial senescence analysis***

To estimate the endothelial cell senescence, we used the average expression levels (Seurat “AddModuleScore”) for the well-established senescence-associated gene sets, including SASP^108,109^ (referred to as EC_SASP), SenMayo^110^, and Reactome pathway R-HSA-2559582^111^), and GO:0090398^112^.

***Pseudotime tracing analysis***

To trace transcriptional evolution of senescent endothelial cells, we constructed a pseudotime trajectory from venous to senescent venous endothelial populations. CCM venous endothelial cells were normalized, scaled, and analyzed using highly variable genes intersected with senescence-related gene sets. Harmony was used for batch correction, Monocle3 was used for trajectory inference, and genes with q_value < 0.05 and num_cells_expressed > 0 were considered dynamic genes. GO terms and signature scores were used to interpret extracellular matrix organization, inflammatory response, and chemokine activity along pseudotime.

**Mouse single-cell RNA-seq data processing and analysis**

***Tissue dissociation and cell purification***

For mouse whole-brain scRNA-seq, Cdh5-CreERT2; Ccm2fl/fl (Ccm2iECKO; n = 3) and wild-type (WT; n = 3) mice were anesthetized and perfused with cold PBS. Brain tissues were mechanically and enzymatically dissociated, filtered through a 40 μm nylon cell strainer, cleared of red blood cells, washed in DPBS with 2% FBS, and processed for scRNA-seq. For endothelial-cell enrichment, dissociated brain cells were labeled with PE anti-mouse CD31 and FITC anti-mouse CD45, and CD31⁺CD45⁻ cells were sorted as murine brain endothelial cells before scRNA-seq.

***scRNA-seq data analysis and cell type annotation***

Following the same library preparation and sequencing procedure of human samples for single-cell RNA-seq. We analyzed scRNA-seq data from mouse whole brain and FACS-sorted endothelial cells using CellRanger (v7.0.1, mm10). Each dataset was processed separately. Quality control was performed by removing cells with total UMI counts ≤ 500 or ≥ 20,000, genes detected ≤ 200 or ≥ 5,000, or mitochondrial UMI content ≥ 20%. Genes detected in fewer than 5 cells were excluded from downstream analysis. Normalization and scaling were performed following the same protocol used for the human scRNA-seq data. Batch correction was carried out using Harmony. Cell clustering was performed using Seurat “FindNeighbors” (k = 20) and “FindClusters” (resolution = 0.5). Whole-brain clusters were annotated by canonical cell type markers; endothelial subclusters were labeled according to marker genes derived from a large-scale murine endothelial single-cell atlas 85.

***Endothelial senescence analysis***

To enable senescence scoring in mouse endothelial cells, we first converted human senescence gene sets into their mouse aliases via the biomaRt R package (v2.54.1). These gene sets were subsequently used to derive module scores using Seurat “AddModuleScore” function.

**Spatial sequencing data processing and analysis**

***Spatial sequencing sample collection and library preparation***

Formalin-fixed, paraffin-embedded (FFPE) samples passing the RNA quality control (DV200 > 30%) were used for spatial transcriptomic construction and sequencing. Five μm thick sections were mounted onto a Visium HD Gene Expression slide (10x Genomics), baked at 42 °C for 3 h, and dried in a desiccator at room temperature overnight. For deparaffinization, the slide was incubated at 60 °C for 2 h, immersed in xylene, and rehydrated in an ethanol gradient. H-E staining was then performed using Mayer’s hematoxylin (Agilent, 011124), bluing reagent (Agilent, CS70230-2), and alcoholic eosin (Abcam, AB246824). Stained slides were scanned under Aperio CS2 (Leica, Germany), followed by decrosslinking using 0.1N HCl and TE Buffer (pH 9.0) to release RNA that was sequestered by formalin. The stained slide incubated with human whole transcriptome probe panel then transferred to Cytassist (10x Genomics).

Human whole transcriptome probe panel (10×) that consisted of three pairs of specific probes (5’ containing Small RNA Read 2S and 3’ containing poly-A) for mostly gene was hybridized to RNA. Probe pairs were then ligated to seal the junctions between them and to form the single-stranded ligation products. The samples were treated with RNase and permeabilized to release the ligation products. Poly-A portion of the products was then captured by the poly (dT) regions of the capture probes precoated on the Visium slide that also include an Illumina Read 1, spatial barcode, and unique molecular identifier (UMI). Probes were extended to produce spatially barcoded ligated probe products and released from the slide for indexing via Sample Index PCR and final library construction and sequencing. Visium Spatial Gene Expression libraries consisted of Illuminapaired-end sequences flanked with P5/P7. The 16-bp Spatial Barcode and 12-bp UMI were encoded in Read 1, while Read 2S was used to sequence the ligated probe insert.

***Alignment and cell segmentation of Visium HD data***

We processed Visium HD datasets using SpaceRanger (v4.0.1; 10x Genomics) with the GRCh38 human reference genome to generate spatially resolved feature–barcode matrices and associated spatial coordinates. Cell segmentation on H&E images was performed by the StarDist-based nucleus detection model integrated in the spaceranger count workflow. Following nucleus detection, each 2-μm Visium HD square was assigned to its nearest nucleus within a maximum radius specified by the --nucleus-expansion-distance-micron parameter (set to 8 μm). Squares associated with the same nucleus were subsequently aggregated to produce cell-resolved spatial transcriptomes.

***QC and unbiased clustering***

We conducted quality control and downstream analyses using Seurat. Cells were excluded if they met any of the following criteria: (1) UMI counts ≤ 50; (2) expressed ≤ 30 genes; (3) ≥ 20% of mitochondrial UMIs; or (4) ≥ 5% of hemoglobin UMIs. Genes detected in fewer than 10 cells were also removed. The resulting filtered expression matrix was normalized to account for library size and log-transformed using the “NormalizeData” function. Gene expression values were then scaled using “ScaleData”. Principal component analysis was performed on the top 2,000 highly variable genes identified via “FindVariableFeatures” with the ‘vst’ selection method. The first 20 principal components were used as input for UMAP dimensionality reduction via the “RunUMAP” function. Batch effects across samples were corrected using Harmony integration. Cell clustering was performed using the Louvain algorithm, with neighborhood graphs constructed using “FindNeighbors” (k = 20) and clustering performed using “FindClusters” at a resolution of 1. Differentially expressed genes were identified with the “FindAllMarkers” function, and major cell types were annotated based on significant cluster-specific gene markers.

***Spatial association between IGHG⁺ plasma cells and endothelial senescence***

The single-cell spatial transcriptomic data with cell type annotations were imported into scanpy (v1.9.1) 75 for downstream analysis. IGHG⁺ plasma cells were defined as those expressing IGHG1–4 genes. For each endothelial cell, we counted the number of IGHG⁺ plasma cells within 30 μm using a k-d tree–based spatial search (scipy. spatial.cKDTree, scipy v1.7.3) 76. The relationship between IGHG⁺ plasma cell proximity and EC_SASP scores was examined using LOESS regression.

***Correlation analysis between endothelial senescence and signaling pathway activity***

To explore how endothelial senescence might be linked to mTORC1 and NF-κB signaling, we summarized EC_SASP, mTORC1, and NF-κB signature scores at the sample level by averaging across endothelial cells. Pearson correlation coefficients were calculated between EC_SASP and each pathway, and regression lines with 95% confidence intervals were visualized.

***Bulk RNA sequencing***

TPM-normalized bulk RNA-seq data from control and CCM lesion samples were retrieved from three published cohorts: Subhash et al., Lyne et al., and Koskimäki et al. Each dataset was processed separately using log₂ transformation, quantile normalization, and z-score standardization. Correlation analyses were performed between lesion size and plasma cell/IgG signature scores when clinical data were available.

**Prussian blue staining**

Prussian blue staining was performed using a Prussian Blue Iron Stain Kit with Nuclear Fast Red counterstaining (G1422, Solarbio, Beijing, China) according to the manufacturer’s instructions. Briefly, paraffin sections were deparaffinized, rehydrated, incubated with freshly prepared Perls working solution at room temperature, counterstained with nuclear fast red, dehydrated, cleared, and mounted. Ferric iron/hemosiderin-positive deposits were identified as blue staining.

**Immunohistochemistry (IHC) staining**

Immunohistochemistry was performed on FFPE sections. Slides were dewaxed, rehydrated, and subjected to heat-induced antigen retrieval using citrate pH 6 or EDTA pH 9 retrieval buffer. Primary antibodies were incubated for 120 min, followed by secondary-antibody amplification according to the manufacturer’s instructions. Immunoreactive cells were visualized with DAB and counterstained with hematoxylin. Images were acquired using a Zeiss Axio Scope A1 microscope and quantified using ImageJ.

**Multiple immunohistochemistry (mIHC) staining**

Multiplexed IHC was performed on FFPE sections using the Opal Multiplex IHC Assay Kit according to the manufacturer’s instructions. Images were acquired using PhenoImager HT or Zeiss LSM 880 systems and quantified using ImageJ.

**Cell Culture**

Primary human umbilical vein endothelial cells (HUVECs) and Human Brain Microvascular Endothelial Cells (HBMECs) were purchased from ScienCell and cultured in endothelial cell medium (#1001, ScienCell) at 37 °C with 5% CO₂. The mouse brain endothelial cell line bEnd.3 was purchased from the American Type Culture Collection (ATCC, CRL-2299) and cultured according to the supplier’s recommendations. HUVECs were treated with purified human IgG at 1 mg/mL for 24 h; PBS was used as the vehicle control. bEnd.3 cells were treated with purified mouse native IgG at 1 mg/mL for 24 h, with PBS as vehicle control. For pathway inhibition, rapamycin was used at 100 nM and PDTC at 10 μM during IgG stimulation. To inhibit lysosomal acidification and autophagic degradation, cells were treated with bafilomycin A1 (BafA1; 10 nM) for 24 h. Rapamycin (S1039), pyrrolidinedithiocarbamate ammonium (PDTC; S3633), and bafilomycin A1 (BafA1; S1413) were purchased from Selleck Chemicals. For combined knockdown and treatment experiments, cells were transfected with the indicated siRNA, cultured for 24 h, and then treated with IgG, inhibitors, or vehicle for an additional 24 h before analysis.

**Short interfering RNA (siRNA) transfection**

HUVECs were seeded into 6-well plates and transfected at approximately 80% confluence using Lipofectamine 3000 (Cat# L3000015, Invitrogen) according to the manufacturer’s protocol. Cells were transfected with siRNAs targeting CCM2, or with negative-control siRNA, at a final concentration of 50 nM in Opti-MEM.

siCCM2 5’- CAUAGACAAUGCAAAGAGATT -3’

siCCM2 5’- UCUCUUUGCAUUGUCUAUGTT -3’

**SPiDER-β-Galactosidase staining**

OCT-embedded tissue sections were thawed at room temperature, fixed with 4% PFA for 20 min, washed with PBS, and incubated with SPiDER-β-Gal staining solution (SG02, Dojindo) at 4 °C overnight. Images were acquired using a ZEISS LSM 880 confocal microscope and quantified using ImageJ. For co-staining, immunofluorescence was performed after SPiDER-β-Gal staining as previously described.

**Immunofluorescence (IF) staining**

Adherent endothelial cells were seeded on ibidi μ-Slide 8 Well glass-bottom chambers and cultured to 70–80% confluence. Cells were fixed with 4% PFA, permeabilized with 0.3% Triton X-100, blocked with normal goat serum, incubated with primary antibodies overnight at 4 °C, and incubated with fluorophore-conjugated secondary antibodies for 1 h at room temperature. Nuclei were counterstained with DAPI. Images were acquired using a ZEISS LSM 880 confocal microscope and quantified using ImageJ. P21⁺ cells were defined by nuclear P21 signal colocalized with DAPI. ZO-1 intensity was measured at cell-cell junctions, and IgG accumulation was quantified within LAMP1⁺ regions using a thresholded LAMP1 mask.

**Senescence-associated β-Galactosidase (SA-β-Gal) staining**

SA-β-Gal staining of adherent cells was performed using the Cellular Senescence Detection Kit (C0602, Beyotime) according to the manufacturer’s instructions. Cells were fixed for 10 min and incubated with staining solution at 37 °C overnight. Images were captured using a Leica microscope. SA-β-Gal-positive cells were quantified using ImageJ with color deconvolution and thresholding.

**Western blot**

Whole-cell lysates were prepared using RIPA buffer (89901, Thermo). Equal amounts of total protein were loaded for western blot. Electrophoresis was performed at 80V until the samples reached the appropriate position. For membrane transfer, 10× transfer buffer (D1060, Solarbio) was prepared according to the manufacturer's instructions, and the transfer was conducted at 90V for 90 minutes. The membrane was blocked with 5% Bovine Serum Albumin (BSA, A8022, Solarbio). The primary antibody was incubated overnight at the recommended concentration, followed by a 1-hour incubation with the secondary antibody at room temperature. If membrane stripping is required, the membrane was treated with stripping buffer (SW3020, Solarbio) for 20 minutes after exposure, then washed and re-incubated with the primary antibody. The protein bands were visualized using the Immobilon Western Chemiluminescent HRP Substrate (MERCK, WBKLS0500) and imaged with the MiniChemi System (SINSAGE, China). Exposure conditions were optimized to prevent signal saturation. The optical density of the bands was quantified using ImageJ software. For each band, the background intensity was subtracted. The relative protein expression was determined by normalizing the intensity of the target protein to that of GAPDH.

**RNA Extraction and Quantitative Real-Time PCR (qRT-PCR)**

Total RNA was extracted from adherent endothelial cells using an RNA extraction kit (Qiagen, Cat#74104), and cDNA was synthesized from 1 μg total RNA using PrimeScript RT Reagent Kit with gDNA Eraser (#RR047A, Takara). qRT-PCR was performed using TB Green Premix Ex Taq II (#RR082A, Takara). Gene expression was normalized to GAPDH and calculated using the 2⁻ΔΔCt method.

**Isolation and culture of primary human endothelial cells**

Fresh surgical specimens were obtained from patients with CCM, AVMs, or focal epilepsy. Non-lesional cortical tissues removed during epilepsy surgery were used as neurological controls. Samples were transferred immediately into ice-cold sterile PBS after surgical resection. After removal of visible blood clots and necrotic tissue, specimens were washed with PBS, mechanically minced into approximately 2mm³ fragments, and enzymatically dissociated using collagenase type IV (Biosharp, BS165) at 37°C for 30 min with gentle agitation. Digestion was terminated by adding ECM, and the resulting cell suspension was passed through a 70-μm cell strainer.

Endothelial cells were enriched by positive selection for CD31 using CD31 Microbeads (Miltenyi Biotec, 130-091-935) according to the manufacturer’s instructions. The isolated cells were seeded onto glass coverslips and maintained in complete endothelial cell growth medium at 37°C in a humidified atmosphere containing 5% CO₂. All cell preparations were fixed and subjected to immunofluorescence staining 16 hours after plating.

**Lysosomal pH was measuring**

Lysosomal pH was measured using LysoSensor Yellow/Blue DND-160. Cells were incubated with 2 μM DND-160 for 3 min and imaged by confocal microscopy. For calibration, cells were incubated with pH calibration buffers adjusted to pH 4.0, 4.5, 5.0, 5.5, and 6.0 in the presence of nigericin. Fluorescence intensities from the two emission channels were quantified using ImageJ, and the fluorescence ratio was calculated. A standard curve was generated by plotting the fluorescence ratio against the corresponding pH values, and lysosomal pH in experimental samples was calculated from the fitted calibration curve.

**Transmission Electron Microscopy**

Freshly resected human CCM specimens were immersion-fixed in 3% glutaraldehyde for 2 h, washed with PBS, post-fixed in 1% osmium tetroxide for 1 h, dehydrated through graded ethanol and propylene oxide, and embedded in Epon812 resin. Ultrathin sections (100 nm) were stained with 1% uranyl acetate followed by 1% lead citrate, dried overnight, and observed using a transmission electron microscope (HITACHI, Tokyo, Japan) at 1.5k magnification.

**Ultrastructural Identification**

The identification of endothelial cells and lysosomes was performed based on established ultrastructural diagnostic criteria defined in prior studies. The pathological endothelial cells were defined as: a markedly attenuated and flattened morphology lining dilated vascular channels; loss of intercellular junctional complexes; and association with abnormal basement membrane ^113,114^. Lysosomes within the identified endothelial cells were classified into two categories based on their ultrastructural morphology: Primary lysosomes, which appeared as round or oval, membrane-bound vesicles of relatively uniform size and contained homogeneous, electron-dense material; and secondary lysosomes, which presented as irregularly shaped, membrane-bound vesicles enclosing heterogeneous, electron-dense components ^115^. For quantitative analysis, 5 fields of view were randomly selected at 1.5k magnification. In each field, the numbers of primary lysosomes, secondary lysosomes, and total lysosomes within identified endothelial cells were counted. The average counts from the five fields were then calculated.

**In Vivo Therapeutic Interventions**

For in vivo therapeutic studies, Ccm2iECKO mice with MRI-confirmed lesions were randomized according to baseline lesion burden. For CD38 targeting, mice received anti-mouse CD38 monoclonal antibody (Bio X Cell, Catalog #CP052) or mouse IgG2a isotype control (Bio X Cell, Catalog #BE0085) by tail-vein injection at 10 mg/kg twice weekly for 4 weeks. For BCMA antibody treatment, *Ccm2*^iECKO^ mice with MRI-confirmed lesions received anti-mouse BCMA monoclonal antibody or matched isotype control by tail-vein injection at 10 mg/kg twice weekly for 4 weeks. For rapamycin treatment, mice received daily intraperitoneal rapamycin (400 μg per mouse) or vehicle control. In the IgG-rescue group, mouse native IgG (Abcam, ab198772) was initiated concurrently with the first anti-BCMA dose and administered by tail-vein injection at 500 μg per mouse per injection, twice weekly for 4 weeks. Native IgG and anti-BCMA antibody were administered during the same treatment sessions throughout the intervention period. The primary outcome was CCM lesion burden quantified by MRI index; secondary outcomes included survival, Prussian blue staining, IgG deposition, endothelial senescence, and junctional integrity.

**Magnetic resonance imaging (MRI)**

MRI was performed using a 7.0 T small-animal MRI scanner (Bruker, Germany) to evaluate CCM lesions with the following parameters: repetition time (TR) = 600 ms, echo time (TE) = 8 ms, field of view (FOV) = 25 × 25 mm², matrix = 256 × 192, 25 slices, and slice thickness = 0.5 mm. Lesion area and whole-brain area were quantified using ImageJ. MRI index was calculated as total lesion area across 25 images divided by total brain area across 25 images.

**Statistical analyses**

Data analyses were performed in R (v4.2.2), Python (v3.9.18), and GraphPad Prism (10.6.0). Normality was assessed using the Shapiro-Wilk test. Two-group comparisons used unpaired t-test, paired t-test, Wilcoxon rank-sum test, or log-rank test, as appropriate. Multiple-group comparisons used one-way ANOVA or Kruskal-Wallis test, and categorical data were analyzed using the proportion test. Correlations were assessed using Pearson or Spearman correlation. Quantitative data are presented as mean ± SEM unless otherwise indicated. Two-sided p < 0.05 was considered statistically significant. Exact sample sizes, statistical tests, and p values are specified in the figures or figure legends.
